# Body Maps of Sound Pitch and its individual differences in Alexithymia and Depression

**DOI:** 10.1101/2023.07.12.548627

**Authors:** Tatsuya Daikoku, Takato Horii, Shigeto Yamawaki

## Abstract

Sound perception extends beyond the boundaries of auditory sensation, encompassing a profound engagement with the entire human body. This intricate interplay between sound and body sensation has long captivated the interest of researchers. In this study, we examined the relationship between our perception of sound pitch and our bodily position senses, while also exploring the role of emotions in shaping this intriguing cross-modal correspondence. We also compare the topography of pitch-triggered body sensations between depressive, alexithymia, and the control groups, and examine their associations with anxiety. Our findings reveal that individuals with depression and alexithymia experience diffuse and less localized body sensations in response to sound pitch, accompanied by heightened feelings of anxiety and negative emotions. These findings imply that diffuse bodily sensations in response to sound may trigger negative emotions such as anxiety and indicate that monitoring pitch-triggered body sensations could serve as a valuable biomarker for emotional disorders. Our study sheds light on the profound importance of body sense awareness in response to sounds, a phenomenon that may be mediated by interoception. This research enhances our understanding of the intricate relationship between sound, emotions, and the human body, offering insights for potential interventions in emotional disorders.

## 1. Introduction

In the intricate interplay of our multisensory experience, where various senses harmonize to shape our perception of reality, the astounding relationship between auditory stimuli and our physical senses has long enthralled researchers. As our senses harmonize to construct our perceptual experience, the enigmatic connection between the pitch of sound and our vivid bodily sense of position remains a captivating puzzle (Jousmäki et al., 1998; Tajadura-Jiménez et al., 2012). For example, even individuals without auditory-tactile synesthesia (Beauchamp et al., 2008) have been shown to spontaneously associate higher-pitched sounds with sensations of upper, smaller, harder, colder, and drier objects compared to lower-pitched sounds (Walker et al., 2010; Spence, 2020; Eitan et al., 2010). A neural study has provided evidence for anatomical connections between the neural substrates of the primary auditory cortex and those involved in bodily sensation, such as the primary and secondary somatosensory regions (Ro et al., 2013). These findings imply that sound perception can elicit not only auditory experiences but also bodily sensations and that there may be neural mechanisms underlying this crossmodal interaction.

Recent research has proposed that the cross-modal correspondences between pitch and position sense are mediated by emotional factors (Spence, 2020; Sievers et al., 2013). Specifically, different pitches can elicit not only body sensations but also various emotions, which in turn may induce localized body sensations. For instance, a previous study reported that high-pitched sounds and ascending tones tend to be rated as conveying emotions such as happiness, brightness, and speed, in contrast to lower-pitched sounds and descending tones (Collier et al., 1998). However, other research has suggested that high-pitched sounds are not always positive, as screeching and fear-inducing sounds are often high-pitched (Jaquet et al., 2014; Ilie et al., 2006).

Research has shown that emotions can be monitored by the topography of emotion-triggered body sensations (Nummenmaa et al., 2014). Specifically, negative emotions such as fear, anger, sadness, and anxiety tend to elicit localized activation of the upper side of the body, while positive emotions such as happiness and love activate a broader range of the body. This monitoring of emotion-triggered body sensation provides a unique tool for understanding individual differences in emotion representation. For example, emotion-triggered body sensations are weaker in individuals with depression, particularly in response to sadness and fear (Lyons et al., 2021), and weaker and less localized in individuals with alexithymia (Lloyd et al., 2021), which shows the difficulty in identifying and labelling emotions (Taylor et al., 1984).

According to previous studies, individuals with alexithymia and depression, which often co-occurs together with anxiety (Hendryx et al., 1991; Li et al., 2015), have been found to exhibit impaired auditory emotion recognition (Wang et al., 2021) and reduced ability to identify pitch, respectively (Schwenzer et al., 2012). Hence, it is possible that alterations in auditory processing and emotional representation in individuals with alexithymia and depression influence the mapping of sounds to the body through emotions. That is, pitch perception is “emotionally” mediated by body sensation, and monitoring of emotion-triggered body sensation may provide a biomarker for emotional disorders such as depression and alexithymia. However, to the best of our knowledge, few studies have investigated the direct relationships among pitch perception, emotion, and body sensation.

The current study aims to examine the connections between pitch, emotion, and body sensation, Specifically, we investigate the topography of pitch-triggered body sensation and compare it with that of emotion-triggered body sensation as demonstrated in previous research (Nummenmaa et al., 2014). We also compare the topography of pitch-triggered body sensations between depressive, alexithymia, and the control groups based on questionnaires, and examine the relationships with anxiety. We hypothesized that, similar to emotion-triggered body sensations, individuals with depression and alexithymia will exhibit less-localized body sensations to pitch (Lyons et al., 2021; Lloyd et al., 2021). Furthermore, it is possible that less-localized body sensations may be associated with negative emotions such as anxiety.

## 2. Results

The present study consists of body-mapping tests and the following emotional judgements in every 16 sounds (see the STAR Methods section for details). Before each experiment, all participants (*N* = 522) completed the Quick Inventory of Depressive Symptomatology (QIDS-J: Rush et al., 2003; Fujisawa et al., 2010) and the Test of Toronto Alexithymia Scale-20 (TAS-20: Bagby et al., 1994). The participants were then provided with the 16 pure tones with different pitches (A1, D2, G2, C3, F3, A#3, D#4, G#4, C#5, F#5, B5, E6, A6, D7, G7, and C8) with random order. After listening to each of the 16 sounds, the participants were asked to respond with clicks to the position in the body that they felt from the sound, using the body image presented on the screen. Two types of emotional judgements were adopted. The first comprised multiple-choice categorical judgements; that is, in each sound, participants were required to select the best 5 emotional categories in ranking elicited by each sound from a list of 33 categories. The second kind comprised nine-point Likert scales of valence and arousal. We compared the topography of pitch and the corresponding emotional responses among depression, alexithymia, and control groups based on the scores of QIDS-J and TAS-20, respectively (see Table 1). The participants also performed pitch discrimination tests to examine whether the findings of the body map test and the underlying emotions are associated with capacities of pitch discrimination rather than the bodily sensations evoked by a pitch and the emotions that mediate them.

**Table 1.** Demographic and clinical characteristics in Alexithymia and Depression groups.

| **Characteristic** | **Total Sample** | **Non-alexithymia (<45)** | **Alexithymia (>63)** | **p value** |
| --- | --- | --- | --- | --- |
| **N** | 522 | 107 | 102 | - |
| **Age** | 38.8(±0.6) | 42.4(±1.2) | 36.1(±1.3) | <.001 |
| **Sex (% female)** | 61.5 | 69.2 | 58.8 | - |
| **QUIDS-J Score** | 4.7(±0.2) | 2.4(±0.3) | 9.1(±0.5) | <.001 |
| **TAS-20 Total** | 54.5(±0.4) | 40.3(±0.4) | 68.4(±0.4) | <.001 |
| **DIF** | 16.9(±0.3) | 9.8(±0.3) | 24.4(±0.4) | <.001 |
| **DDF** | 15.6(±0.2) | 11.6(±0.3) | 19.8(±0.2) | <.001 |
| **EOT** | 22.0(±0.16) | 18.9(±0.4) | 24.2(±0.3) | <.001 |
| **Characteristic** | **Total Sample** | **Non-depression (=0)** | **Depression (>9)** | **p value** |
| **N** | 522 | 107 | 103 | - |
| **Age** | 38.8(±0.6) | 35.7(±1.3) | 37.1(±1.3) | 0.45 |
| **Sex (% female)** | 61.5 | 59.8 | 64.1 | - |
| **QUIDS-J Score** | 4.7(±0.2) | 12.7(±0.3) | 0(±0) | <.001 |
| **TAS-20 Total** | 54.5(±0.4) | 50.5(0.9) | 62.9(±1.0) | <.001 |
| **DIF** | 16.9(±0.3) | 14.9(±0.5) | 22.1(±0.6) | <.001 |
| **DDF** | 15.6(±0.2) | 13.5(±0.3) | 18.0(±0.4) | <.001 |
| **EOT** | 22.0(±0.16) | 22.1(±0.3) | 22.8(±0.4) | 0.2 |

### 2.1. Body map

Grand averages of pitch-triggered topography and the clicked positions were shown in supplementary materials (S1 and S2 Appendix). All of the anonymized raw data files and all of the results of statistical analyses including the descriptives have been deposited to an external source (https://osf.io/rs4kh/?view_only=cbe06bcaac4d42dc809f1a54cb173596).

The individuals with alexithymia (Figure 1) and depression (Figure 2) showed less-localised and diffused body sensations to pitch. Statistical analysis indicated that the number of different click positions and the total number of clicks is increased in individuals with Alexithymia and Depression groups. The one-way analysis of variance (ANOVA) detected that the number of different clicks positions is significantly higher in individuals with alexithymia compared to the control, specifically in response to the sounds of D#4 (χ*² =* 3.89, *p =* .049, ε*² =* .02) and C#5 (χ*² =* 3.86, *p =* .049, ε*² =* .02). The number of different clicks positions is significantly higher in individuals with depression compared to the control, specifically in response to the lower-pitched sounds of A1 (*χ² =* 7.01, *p =* .008, *ε² =* .03), D2 (*χ² =* 7.14, *p =* .008, *ε² =* .03), G2 (*χ² =* 9.28, *p =* .002, *ε² =* .04), C3 (*χ² =* 7.18, *p =* .007, *ε² =* .03), F3 (*χ² =* 9.95, *p =* .002, *ε² =* .05), A#3 (*χ² =* 6.52, *p =* .01, *ε² =* .03), and D#4 (*χ² =* 6.60, *p =* .01, *ε² =* .03).

**Figure 1.**
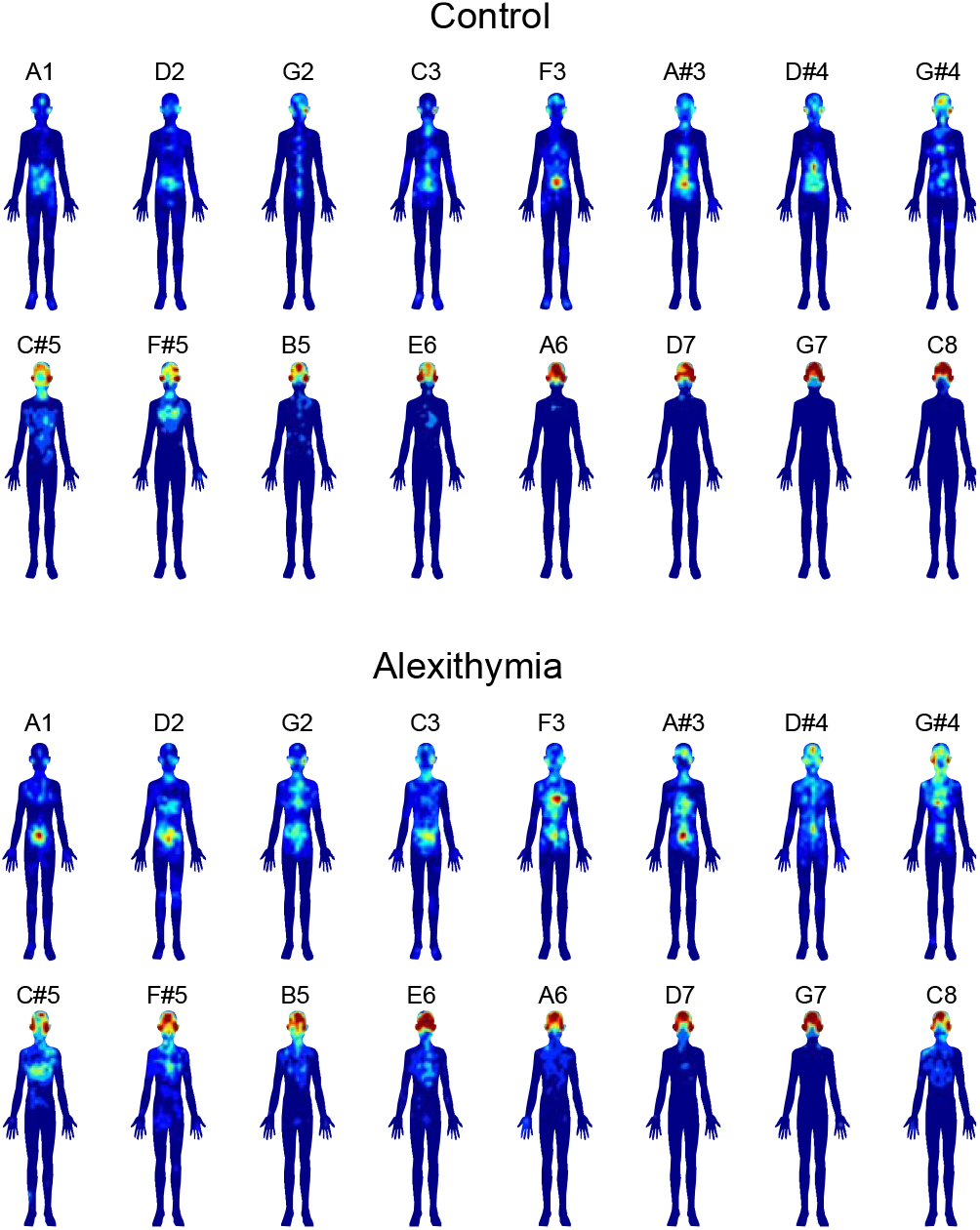
Body topography of pitches in the Alexithymia group. The blue-to-red gradients represent the number of clicks. Compared to the control group, individuals with Alexithymia exhibit a more diffuse and non-localized body map.

**Figure 2.**
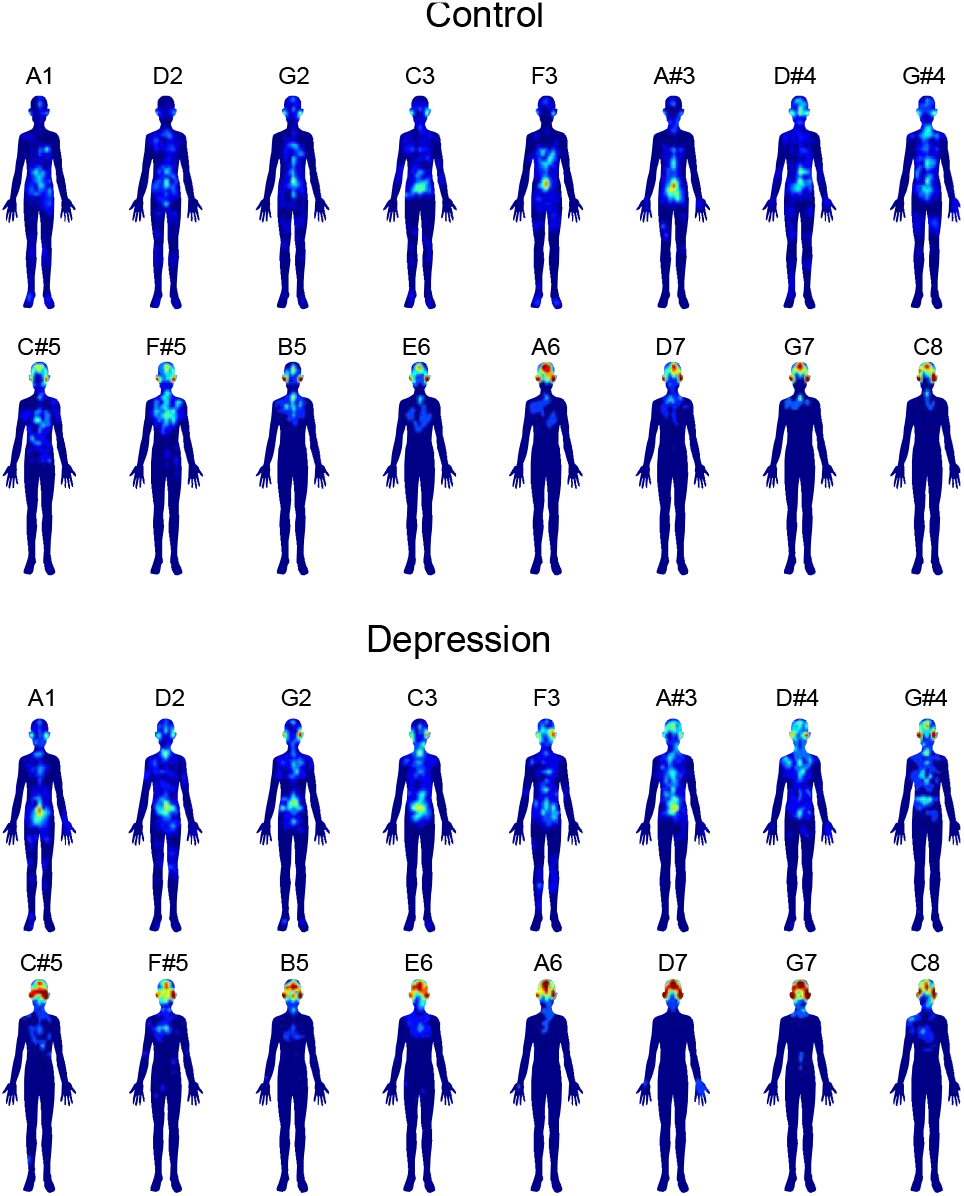
Body topography of pitches in the Depression group. The blue-to-red gradients represent the number of clicks. Compared to the control group, individuals with Depression exhibit a more diffuse and non-localized body map.

The total number of clicks is significantly higher in individuals with depression compared to the control, specifically in response to the lower-pitched sounds of A1 (*χ² =* 7.11, *p =* .008, *ε² =* .03), D2 (*χ² =* 7.39, *p =* .007, *ε² =* .04), G2 (*χ² =* 9.06, *p =* .003, *ε² =* .04), C3 (*χ² =* 7.19, *p =* .007, *ε² =* .03), F3 (*χ² =* 10.04, *p =* .002, *ε² =* .05), A#3 (*χ² =* 6.73, *p =* .009, *ε² =* .03), and D#4 (*χ² =* 6.61, *p =* .01, *ε² =* .03). No statistical difference in the total number of clicks was observed between individuals with alexithymia and the control.

### 2.2. Emotion in response to pitches

All of the results of statistical analyses and the descriptives of the multiple-choice categorical judgements and nine-point Likert scale of valence and arousal have been deposited to an external source. The figures of the grand average data are shown in S3 and S4 Appendices (Figure a) in supplementary materials.

The multiple-choice categorical judgements test showed that the individuals with alexithymia and depression felt strong anxiety in response to several pitched sounds (see Figure b, S4 Appendix). The one-way ANOVA detected that the anxiety is significantly higher in individuals with alexithymia compared to the control, specifically in response to the lower-pitched sounds of A1 (*χ² =* 6.86, *p =* .009, *ε² =* .03), D2 (*χ² =* 6.20, *p =* .01, *ε² =* .03), G2 (*χ² =* 4.52, *p =* .03, *ε² =* .02), A#3 (*χ² =* 5.50, *p =* .02, *ε² =* .03), and D#4 (*χ² =* 17.75, *p <* .001, *ε² =* .09). The anxiety is significantly higher in individuals with depression compared to the control, in response to the wide range of pitched sounds of A1 (*χ² =* 6.03, *p =* .01, *ε² =* .03), D2 (*χ² =* 4.99, *p =* .03, *ε² =* .02), C3 (*χ² =* 4.94, *p =* .03, *ε² =* .02), F3 (*χ² =* 13.02, *p <* .001, *ε² =* .06), D#4 (*χ² =* 7.36, *p =* .007, *ε² =* .04), F#5 (*χ² =* 4.02, *p =* .045, *ε² =* .02), B5 (*χ² =* 6.42, *p =* .011, *ε² =* .03), E6 (*χ² =* 5.48, *p =* .019, *ε² =* .03), A6 (*χ² =* 5.51, *p =* .019, *ε² =* .03), and G7 (*χ² =* 9.70, *p =* .002, *ε² =* .05).

The nine-point Likert scale of valence and arousal detected that the individuals with alexithymia and depression felt negative valence in response to several pitched sounds while no significant differences in arousal between the individuals with alexithymia and the controls and between the individuals with depression and the controls were observed (Figure b, S4 Appendix). The one-way ANOVA detected that the valence is significantly negative in individuals with alexithymia compared to the control, in response to the lower-pitched sounds of D2 (*χ² =* 4.13, *p =* .04, *ε² =* .02), C3 (*χ² =* 4.59, *p =* .03, *ε² =* .02), F3 (*χ² =* 5.05, *p =* .03, *ε² =* .02), and A#3 (*χ² =* 7.04, *p =* .008, *ε² =* .03). The valence is significantly negative in individuals with depression compared to the control, in response to the several pitched sounds of C3 (*χ² =* 4.89, *p =* .03, *ε² =* .02), F3 (*χ² =* 4.37, *p =* .04, *ε² =* .02), and A#3 (*χ² =* 5.40, *p =* .02, *ε² =* .03).

The participants also performed pitch discrimination tests after experiments of body sensation. The results showed no statistical significance between groups (Depression vs. control: p = 0.45; Alexithymia vs. control: p = 0.86, see Figure b in S1 Appendix). This suggests that not pitch discrimination ability per se, but rather the bodily sensations evoked by pitch and the emotions that mediate them may be implicated in modulating pitch-triggered topography.

## 3. Discussion

The present study investigated individual differences in pitch-triggered body sensations between depression, alexithymia, and a control group based on questionnaires (QIDS and TAS-20), and how these differences are associated with emotional responses such as anxiety and valence. The results indicated that alexithymia and depression groups exhibited less localized and more diffuse body sensations in response to pitch while there were no significant group differences in pitch discrimination ability. Furthermore, individuals with alexithymia and depression experienced strong feelings of anxiety and negative valence in response to several pitches. These results suggest that the diffuse bodily sensations are not inherently linked to pitch discrimination ability, but rather to the emotional responses triggered by sound pitches. This study suggests that the diffuse bodily sensations in response to pitch may induce negative emotions such as anxiety.

Previous studies have suggested that emotion-triggered body sensations are modulated in individuals with depression (Lyons et al., 2021) and alexithymia (Lloyd et al., 2021). Specifically, individuals with alexithymia exhibit weaker and less localized emotion-triggered body sensations (Lloyd et al., 2021). Our study extends these findings by demonstrating that pitch-triggered body sensations are also less localized in alexithymia and depression, and that this diffuse sensation is associated with feelings of anxiety. For instance, the pitch of D#4 induces diffuse body sensations and strong anxiety in individuals with alexithymia compared to controls. It is possible that the diffuse body sensation is mediated by anxiety, which is frequently co-occurring with alexithymia and depression (Hendryx et al., 1991; Li et al., 2015). Previous evidence has demonstrated that individuals with alexithymia have impaired auditory emotion recognition (Wang et al., 2021), suggesting that alterations in auditory processing and emotional representation in these individuals may affect the mapping of sounds to the body through emotions. Our findings suggest that pitch perception is “emotionally” mediated by body sensation, and monitoring of emotion-triggered body sensation of sounds may provide a biomarker for emotional disorders such as depression and alexithymia.

Previous studies have examined the relationship between pitch discrimination ability and mood disorders. They found that the pitch discrimination ability is enhanced in depression and that this can also be observed from the mismatch negativity (MMN) using electroencephalography (EEG) (Bonetti et al., 2017), whereas the pitch discrimination ability is decreased in schizophrenia (McLachlan et al., 2013). While our study did not find a significant group difference in pitch discrimination ability, we did observe that individuals with depression perceive pitched sounds differently than those without depression, as evidenced by a significantly higher total number of clicks (Figure b, S4 Appendix).

Our study focused on the position sense regarding pitch, which may be associated with the proprioception of sound, but our results also suggest a possible relationship with interoception for some reasons. The first is that the lowest bodily location felt in response to pitch was not the feet, located at the lowest point of the body, but rather the abdomen, the lowest part of the visceral system. The second is that the body sensation evoked by emotions is suggested to entail interoceptive involvement (Nummenmaa et al., 2014). It is possible that the diffuse body sensation of sounds could reflect a diminished “*bodily or interoceptive awareness*” in response to sound. That is, interoceptive awareness, which involves perceiving and interpreting bodily signals, may be altered in individuals with these emotional disorders. Understanding the role of interoceptive awareness in the perception of sounds and emotions has important implications for the development of interventions for individuals with sensory processing disorders, as well as for alexithymia and depression.

The present study was unable to strictly control the auditory environment due to the online experiment. For instance, it is probable that the negative valence is partially correlated with volume. However, in this study, all participants were instructed to use headphones, which ensured that the volume did not vary drastically among participants, in contrast to using speakers. Another limitation is that grouping was conducted only based on the questionnaire, which may have led to the less strict categorization of alexithymia and depression. However, given the large sample size of over 500 participants in this study, the noise in the responses is likely to have been offset. Given that even when targeting individuals assessed solely through questionnaires for alexithymia and depression, we detected groups difference in body map of sounds. This suggests that the body map of sounds may prove to be a valuable tool in the discovery of latent factors and early diagnosis. Finally, further research is needed to examine how musical education and culture influence the body sensation to sounds and to investigate whether the body sensation to sounds and the associated emotional responses are modulated depending on not only pitch but also differences in timbre or the semantic meaning of sound.

This study may shed light on potential interventions for alexithymia and depression by exploring ways to localize body sensation in response to pitch. Prior research has suggested that music-based interventions may improve emotion recognition in adolescents with alexithymia (Pedregal et al., 2021). Additionally, research has suggested that emotion-triggered body sensations become more localized over development to adult-like patterns (Hietanen et al., 2016). Therefore, understanding what kinds of sounds are ideal for each individual’s development, as well as for alexithymia and depression, is important insight into applying to the potential interventions. Furthermore, by comprehending which types of sound and timbre cause discomfort or anxiety for different personalities or developmental stages, it may be possible to provide an optimal auditory environment for each individual. Thus, further research in this area is crucial.

## 4. Conclusions

The present study suggests that individuals with alexithymia and depression experience diffuse bodily sensations and strong anxiety and negatively-valenced emotions in response to sound pitch. These findings imply that diffuse bodily sensations in response to sound may trigger negative emotions such as anxiety. The present study may suggest the importance of interoceptive awareness in the processing of pitch-triggered body sensations and its association with emotional responses. This study may also provide insight into potential interventions for alexithymia and depression by exploring methods to localize body sensations in response to pitch. Specifically, by understanding which types of sound and timbre cause discomfort or anxiety for different personalities or developmental stages, it may be possible to create an optimal auditory environment for each individual. These findings have important implications for the treatment of individuals with alexithymia and depression and may inform the development of personalized therapeutic interventions.

## 4. Methods

### 4.1. Participants

The present study consists of body-mapping tests and the following emotional judgements in every 16 sounds. The Japanese participants took part in the study (N = 522, *M*_age_ = 38.8, female = 321, see Table 1 for Demographic characteristics). They have no history of neurological or audiological disorders and no absolute pitch. The experiment was conducted in accordance with the guidelines of the Declaration of Helsinki and was approved by the Ethics Committee of The University of Tokyo (Screening number: 21-335). All participants gave their informed consent and conducted the experiments by PC.

Before each experiment, all participants completed the Quick Inventory of Depressive Symptomatology (QIDS-J) (Rush et al., 2003; Fujisawa et al., 2010 for Japanese version) and the Test of Toronto Alexithymia Scale-20 (TAS-20) (Bagby et al., 1994) (see S4 Appendix). In order to balance the number of participants across groups and to compare groups with scores at opposite ends of the spectrum in each test, we divided them into groups with extremely high scores (indicating strong alexithymia or severe depression) and those with extremely low scores on these tests. Demographic and clinical characteristics in Alexithymia and Depression groups are shown in Table 1.

Using the scores of TAS-20, with an aim to achieve a balanced distribution of approximately 100 participants in each group, we divided into two group: a group of individuals who showed higher scores than 63 (hereafter, “*alexithymia group*”) and a group of individuals who showed lower than 45 (hereafter, control group). The TAS-20, a widely recognized measurement tool developed by Bagby et al. (1994), is employed to assess alexithymia, a condition characterized by difficulties in identifying and expressing one’s emotions. Consisting of 20 items, this scale is commonly used in research settings. Respondents rate each item on a 5-point Likert-type scale, ranging from “Strongly Disagree” to “Strongly Agree.” To ensure accuracy, certain items (4, 5, 10, 18, and 19) are reverse scored. In our analysis, we focused on the total score obtained by summing the individual item scores. Previous studies have demonstrated that the TAS-20 possesses satisfactory internal consistency (α = .81) and test-retest reliability (.77, p < .01) (Bagby et al., 1994). Researchers often employ specific cut-off scores to interpret the results. Scores equal to or below 51 indicate the absence of alexithymia, scores falling between 52 and 60 suggest possible alexithymia, and scores equal to or above 61 indicate the presence of alexithymia. During our study, we encountered an issue related to item #19 in the survey questionnaire. This particular item was displayed improperly, leading to a high non-response rate (5.2%) or neutral responses (16.3% rated as “neither agree nor disagree”). To address this concern, we made the decision to exclude item #19 from the analysis and subsequently adjusted the range and cut-off scores accordingly. Following these modifications, the adjusted cut-off scores for the TAS-20 are as follows: total scores equal to or below 46 indicate the absence of alexithymia, scores ranging from 47 to 55 suggest possible alexithymia, and scores equal to or above 56 indicate the presence of alexithymia. By utilizing the TAS-20 total score, which has been adjusted in line with previous research practices (e.g., Starita & di Pellegrino, 2018), we categorized participants into two groups: a non-alexithymia group (TAS-20 total score equal to or below 46) and a probable alexithymia group, representing scores consistent with possible to present alexithymia (TAS-20 total score above 63). These adjustments align with criticisms that the original cut-off score for alexithymia (TAS-20 ≥ 61) is overly conservative (Franz et al., 2008).

Furthermore, using the scores from the QIDS-J, with an aim to achieve a balanced distribution of approximately 100 participants in each group, we categorized individuals into two groups. One group consisted of individuals who had the highest scores (≧ 9, hereafter, “*depression group*”), while the other group consisted of individuals who had the lowest scores (=0, hereafter, control group). The QIDS contain 16 items that measure the nine DSM-IV-TR criterion symptom domains including sad mood, poor concentration, self-criticism, suicidal ideation, anhedonia, energy/fatigue, sleep disturbance, decrease/increase in appetite/weight, and psychomotor agitation/retardation and were designed to measure overall severity of the depressive syndrome. The items related to sleep (Items 1-4), appetite/weight (Items 6-9), and psychomotor status (Items 15 and 16) are scored by selecting the highest score among the items within each category. The remaining items (Items 5, 10, 11, 12, 13, and 14) are scored individually. The severity of depression is evaluated by combining scores from the sleep, appetite/weight, psychomotor, and other 6 items, resulting in a total score ranging from 0 to 27. Since each item corresponds to symptoms of major depressive disorder, this evaluation tool can be used for assessing and screening depressive symptoms. Additionally, by calculating the total score, changes in the depressive state can be observed. The QIDS depression severity thresholds have been suggested of 6 to10 for mild, 11 to 15 for moderate, 16 to 20 for severe, and 21+ for very severe depression. Therefore, the Depressive group in this study includes mild to severe depressive symptoms (Table 1).

### 4.2. Materials and Methods

The experimental paradigm was generated using Gorilla Experiment Builder (https://gorilla.sc), which is a cloud-based research platform that allows deploying of behavioural experiments online. After the questionnaires of TAS-20 and QUIDS-J, each participant was provided with the 16 pure tones (20 seconds, 44.1kHz, 32bit, amplitude based on equal-loudness-level contours, Robinson et al., 1956) with different pitches (A1: 55Hz, D2: 73.4Hz, G2: 98Hz, C3: 130.8Hz, F3: 174.6Hz, A#3: 233.1Hz, D#4: 311.1Hz, G#4: 415.3Hz, C#5: 554.4Hz, F#5: 740Hz, B5: 987.8Hz, E6: 1318.5Hz, A6: 1760.0Hz, D7: 2349.3Hz, G7: 3136Hz, and C8: 4186.0Hz) with random order. Each 16-tone was preceded by 1 second of pure tone (A4: 400 Hz) and 500 ms of silence. All original sound files are available freely from Open Science Repository (https://osf.io/rs4kh/?view_only=cbe06bcaac4d42dc809f1a54cb173596).

After listening to each of the 16 sounds, the participants were asked to respond with clicks to the position in the body that they felt from the sound using the body image presented on the screen. The clicking was allowed any number of times up to a maximum of 100 clicks, and clicking while listening to the sound was also allowed (see Figure a of the S2 Appendix for the details).

Two surveys were used to obtain emotional judgements. The first comprised multiple-choice categorical judgements; that is, in each sound, participants were required to select the best 5 emotional categories in ranking elicited by each sound from a list of 33 categories (see S3 Appendix). The 33 emotion categories were derived from emotion taxonomies of prominent theorists, Keltner and Lerner (2010) and based on the previous study by Cowen et al., (2017; 2019; 2020). The second kind comprised nine-point dimensional judgments; that is, after hearing the sounds, participants were required to rate each sound along the valence and arousal. Each rating was obtained on a nine-point Likert scale with the number 5 anchored at neutral.

After all 16 tones were answered, participants performed 10 types of pitch discrimination tests. After one second of a pure tone at 440 Hz (A4, 44.1kHz, 32bit) followed by 500 ms of silence, one second of pure tone with each 20-cent different pitch (44.1kHz, 32bit) was presented with random order (+20cent: 445.1Hz, +40cent: 450.3Hz, +60cent: 455.5Hz, +80cent: 460.8Hz, +100cent: 466.2Hz, -20cent: 434.9Hz, -40cent: 430.0Hz, -60cent: 425.0Hz, -80cent: 420.1Hz, - 100cent: 415.3Hz). The participants were asked to answer whether the second tone was lower or higher than the first tone (i.e., 440Hz) by forced choice. A training test in which a 440 Hz pitch and 500 ms of silence followed by 660 Hz was conducted before the 10 types of pitch discrimination tests.

### 4.3. Statistical Analysis

Using the coordinate data of x and y in body mapping test, we extracted the number of different click positions and the total number of clicks in each participant. The raw data of x and y coordinates (see Figure b in S2 Appendix) were downsampled by a factor of 40. The Figures of the body topographies of pitches (Figure 1 and 2) were generated using Matlab (2022b) by interpolating the coordinates of x and y in a meshgrid format with a color map that represented the neighboring points. The results of the best 5 emotional categories in the ranking were used to score the intensity of 33 emotions. That is, the first, second, third, fourth, and fifth categories were each scored as a 5, 4, 3, 2, and 1 point. The scores of each 33 emotional categories were then averaged for all participants and each group (see S4 Appendix for all results). The past evidence has shown that individuals with alexithymia and depression often co-occurs together with anxiety (Hendryx et al., 1991; Li et al., 2015) and that the negative emotion modulates the emotion-triggered body sensations in individuals with depression (Lyons et al., 2021). Given these findings, the present study particularly focuses on the emotion of anxiety. Measuring both positive and negative affect can be a reliable method of obtaining responses (Watson, Clark, & Tellegen, 1988). In this study, we utilized 33 items to assess various emotions, including anxiety and its opposite of Japanese-term, relief. To reverse the 1-5 scale to 5-1, we scored relief in the opposite direction. We then calculated the average of the anxiety score and the reversed relief score.

We performed the Shapiro–Wilk test for normality on the number of different click positions, the total number of clicks in each participant, the anxiety score of the multiple-choice categorical judgements, and the valence and arousal scores of the nine-point dimensional judgments. Depending on the result of the test for normality, either the parametric or non-parametric (Kruskal-Wallis) One-Way analysis of variance (ANOVA) was applied to compare Alexithymia vs. control, and Depression vs. control groups in each pitch. We also performed the independent samples T-test for the average score of the 10 types of the pitch discrimination test. Statistical analyses were conducted using jamovi Version 1.2 (The jamovi project, 2021). We selected *p* < .05 as the threshold for statistical significance.

## References

Bagby, M., Parker, J. D. A., & Taylor, G. J. (1994). The twenty-item selection Toronto Alexithymia Scale - I. Item selection and and cross-validation structure. Journal of Psychosomatic Reserach, 38(1), 23–32.

Beauchamp, M. S., & Ro, T. (2008). Neural substrates of sound–touch synesthesia after a thalamic lesion. Journal of Neuroscience, 28(50), 13696–13702.

Collier, W. G., & Hubbard, T. L. (1998). Judgments of happiness, brightness, speed and tempo change of auditory stimuli varying in pitch and tempo. Psychomusicology: A Journal of Research in Music Cognition, 17(1-2), 36.

Cowen, A. S., Fang, X., Sauter, D. A., & Keltner, D. (2020). What music makes us feel: uncovering 13 kinds of emotion evoked by music across cultures. Proc. Natl. Acad. Sci. USA, 117, 1924–1934.

Cowen, A. S., & Keltner, D. (2017). Self-report captures 27 distinct categories of emotion bridged by continuous gradients. Proceedings of the national academy of sciences, 114(38), E7900–E7909.

Cowen, A. S., Elfenbein, H. A., Laukka, P., & Keltner, D. (2019). Mapping 24 emotions conveyed by brief human vocalization. American Psychologist, 74(6), 698.

Eitan, Z., & Timmers, R. (2010). Beethoven’s last piano sonata and those who follow crocodiles: Cross-domain mappings of auditory pitch in a musical context. Cognition, 114(3), 405-422.

Eitan, Z., & Rothschild, I. (2011). How music touches: Musical parameters and listeners’ audio-tactile metaphorical mappings. Psychology of Music, 39(4), 449–467.

Fijisawa D, Nakagawa A, Tajima M et al. Development of own Japanese edition entry system simple depression linear measure (Japanese edition QIDS-SR). Jap. J. Stress Sci. 2010; 25: 43–52.

Hendryx, M. S., Haviland, M. G., & Shaw, D. G. (1991). Dimensions of alexithymia and their relationships to anxiety and depression. Journal of Personality Assessment, 56(2), 227– 237.

Hietanen, J. K., Glerean, E., Hari, R., & Nummenmaa, L. (2016). Bodily maps of emotions across child development. Developmental Science, 19(6), 1111–1118.

Ilie, G., & Thompson, W.F. (2006). A comparison of acoustic cues in music and speech for three dimensions of affect. Music Perception, 23, 319–329.

Jaquet, L., Danuser, B., & Gomez, P. (2014). Music and felt emotions: How systematic pitch level variations affect the experience of pleasantness and arousal. Psychology of Music, 42(1), 51–70.

Jousmäki, V., & Hari, R. (1998). Parchment-skin illusion: sound-biased touch. Current biology, 8(6), R190–R191.

Kantrowitz, J. T., Hoptman, M. J., Leitman, D. I., Moreno-Ortega, M., Lehrfeld, J. M., Dias, E., … & Javitt, D. C. (2015). Neural substrates of auditory emotion recognition deficits in schizophrenia. Journal of Neuroscience, 35(44), 14909–14921.

Keltner, D., & Lerner, J. S. (2010). Emotion. In D. T. Gilbert, S. T. Fiske, & G. Lindzey (Ed.), The Handbook of Social Psychology (pp. 317–352). New York, Wiley.

Koenig, L., & Ro, T. (2022). Sound Frequency Predicts the Bodily Location of Auditory-Induced Tactile Sensations in Synesthetic and Ordinary Perception. bioRxiv.

Ley-Flores, J., Alshami, E., Singh, A., Bevilacqua, F., Bianchi-Berthouze, N., Deroy, O., & Tajadura-Jiménez, A. (2022). Effects of pitch and musical sounds on body-representations when moving with sound. Scientific reports, 12(1), 1–20.

Li, S., Zhang, B., Guo, Y., & Zhang, J. (2015). The association between alexithymia as assessed by the 20-item Toronto Alexithymia Scale and depression: A meta-analysis. Psychiatry Research, 227(1), 1– 9.

Lloyd, C. S., Stafford, E., McKinnon, M. C., Rabellino, D., D’Andrea, W., Densmore, M., … & Lanius, R. A. (2021). Mapping alexithymia: Level of emotional awareness differentiates emotion-specific somatosensory maps. Child Abuse & Neglect, 113, 104919.

Lyons, N., Strasser, A., Beitz, B., Teismann, T., Ostermann, T., Anderle, L., & Michalak, J. (2021). Bodily maps of emotion in major depressive disorder. Cognitive Therapy and Research, 45(3), 508–516.

Pedregal, C. R., & Heaton, P. (2021). Autism, music and Alexithymia: A musical intervention to enhance emotion recognition in adolescents with ASD. Research in Developmental Disabilities, 116, 104040.

Ro, T., Ellmore, T. M., & Beauchamp, M. S. (2013). A neural link between feeling and hearing. Cerebral cortex, 23(7), 1724–1730.

Robinson, D. W., & Dadson, R. S. (1956). A re-determination of the equal-loudness relations for pure tones. British Journal of Applied Physics, 7(5), 166.

Rusconi, E., Kwan, B., Giordano, B. L., Umilta, C., & Butterworth, B. (2006). Spatial representation of pitch height: the SMARC effect. Cognition, 99(2), 113–129.

Rush AJ, Trivedi MH, Ibrahim HM et al. The 16-item Quick Inventory of Depressive Symptomatology (QIDS), Clinician Rating (QIDS-C), and Self-Report (QIDS-SR): A psychometric evaluation in patients with chronic major depression. Biol. Psychiatry 2003; 54: 573–583.

Schwenzer, M., Zattarin, E., Grözinger, M., & Mathiak, K. (2012). Impaired pitch identification as a potential marker for depression. BMC psychiatry, 12, 1–6.

Sievers, B., Polansky, L., Casey, M. & Wheatley, T. Music and movement share a dynamic structure that supports universal expressions of emotion. Proc. Natl. Acad. Sci. 110, 70–75 (2013).

Spence, C. (2020). Simple and complex crossmodal correspondences involving audition. Acoustical Science and Technology, 41(1), 6–12.

Tajadura-Jiménez, A., Väljamäe, A., Toshima, I., Kimura, T., Tsakiris, M., & Kitagawa, N. (2012). Action sounds recalibrate perceived tactile distance. Current Biology, 22(13), R516–R517.

Taylor, G. J. (1984). Alexithymia: concept, measurement, and implications for treatment. The American journal of psychiatry.

Walker, P., Bremner, J. G., Mason, U., Spring, J., Mattock, K., Slater, A., & Johnson, S. P. (2010). Preverbal infants’ sensitivity to synaesthetic cross-modality correspondences. Psychological Science, 21(1), 21–25.

Wang, Z., Chen, M., Goerlich, K. S., Aleman, A., Xu, P., & Luo, Y. (2021). Deficient auditory emotion processing but intact emotional multisensory integration in alexithymia. Psychophysiology, 58(6), e13806.

Watson, D., Clark, L. A., & Tellegen, A. (1988). Development and validation of brief measures of positive and negative affect: the PANAS scales. Journal of personality and social psychology, 54(6), 1063.

