## Supplementary material for "Body Maps of Sound Pitch and its individual differences in Alexithymia and Depression": S1 Appendix

**Figure a. the Body map of tone in each group and each tone.**

**Grand average of all participants**

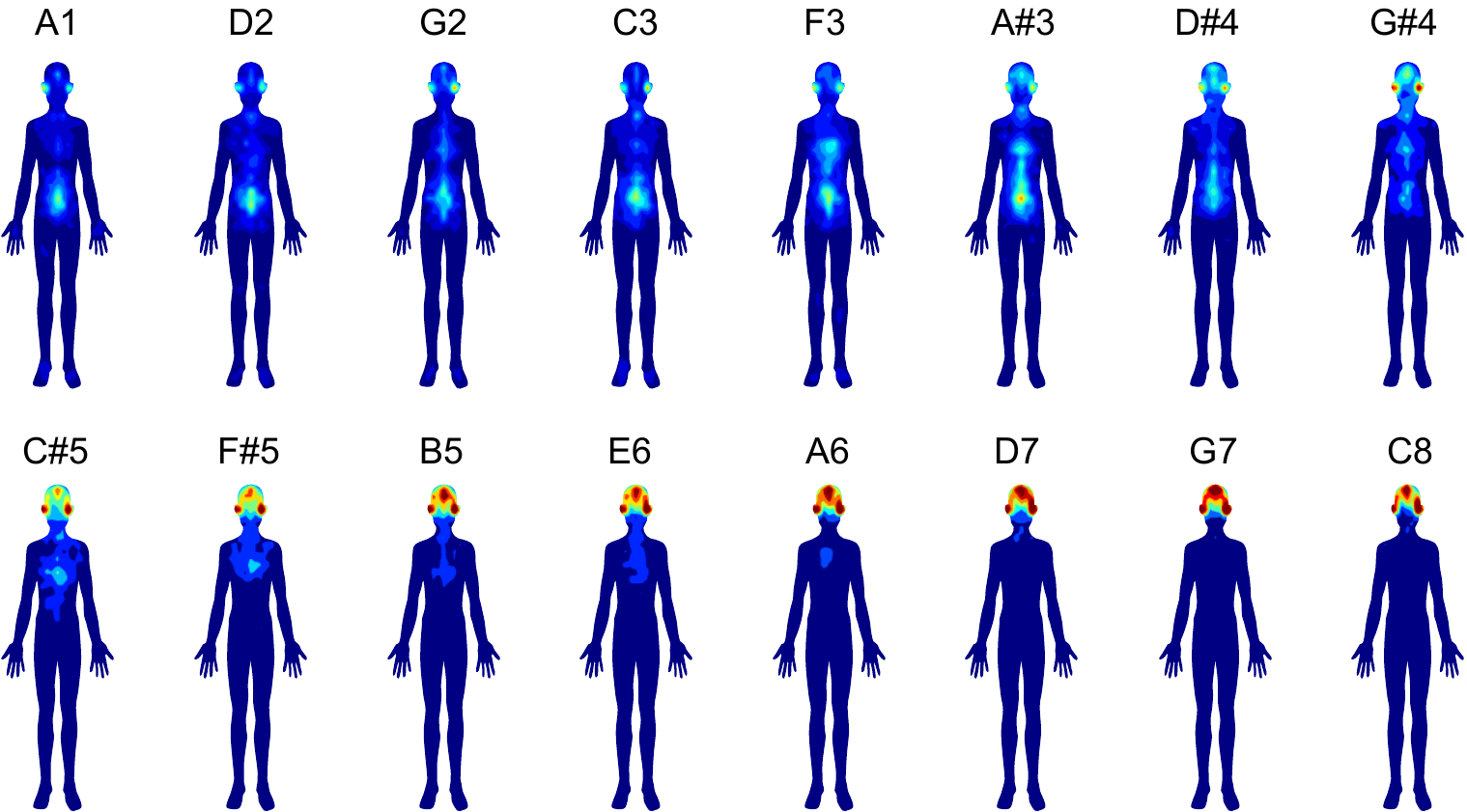

**Depression Group**

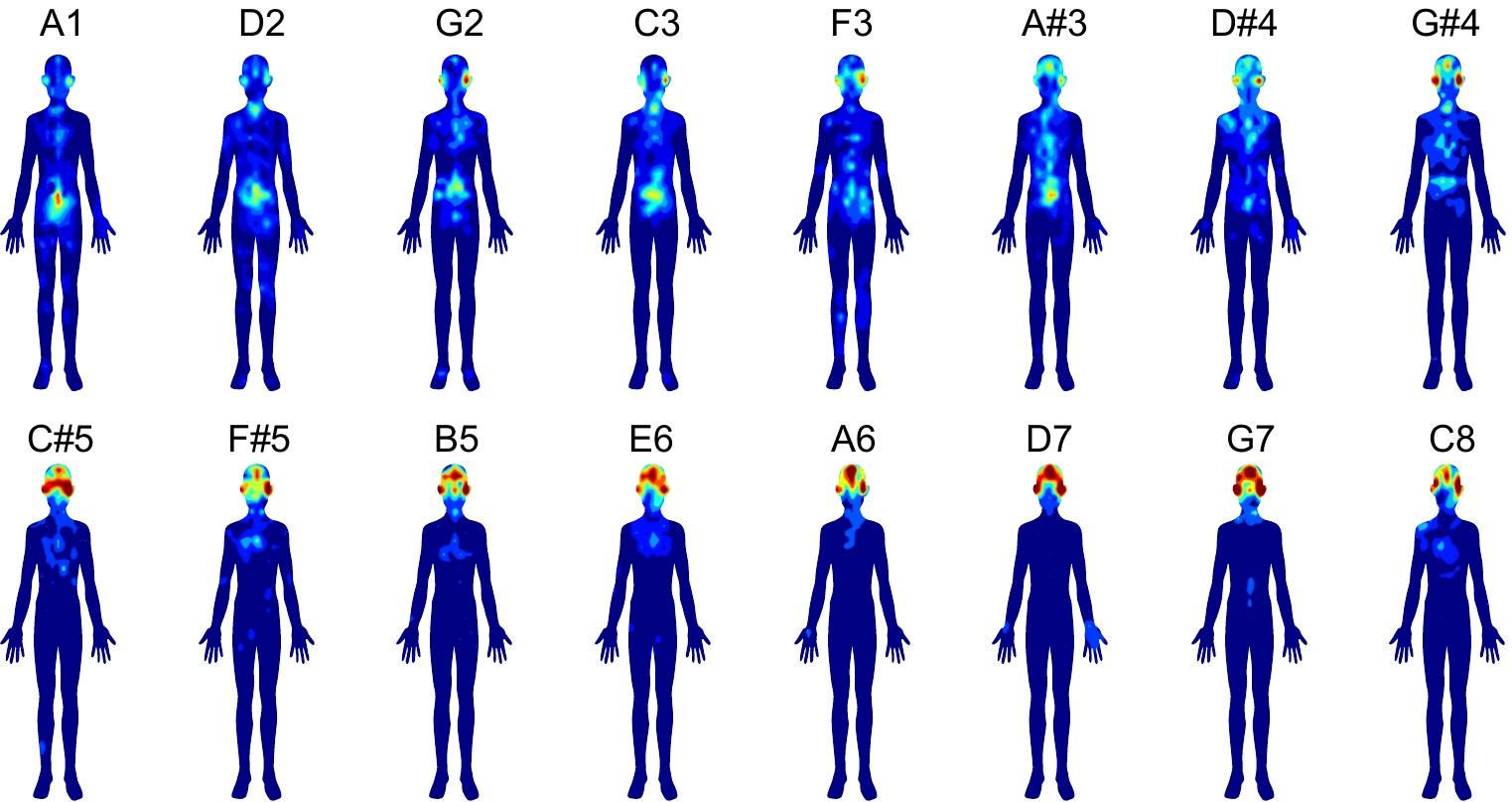

**Non depression Group**

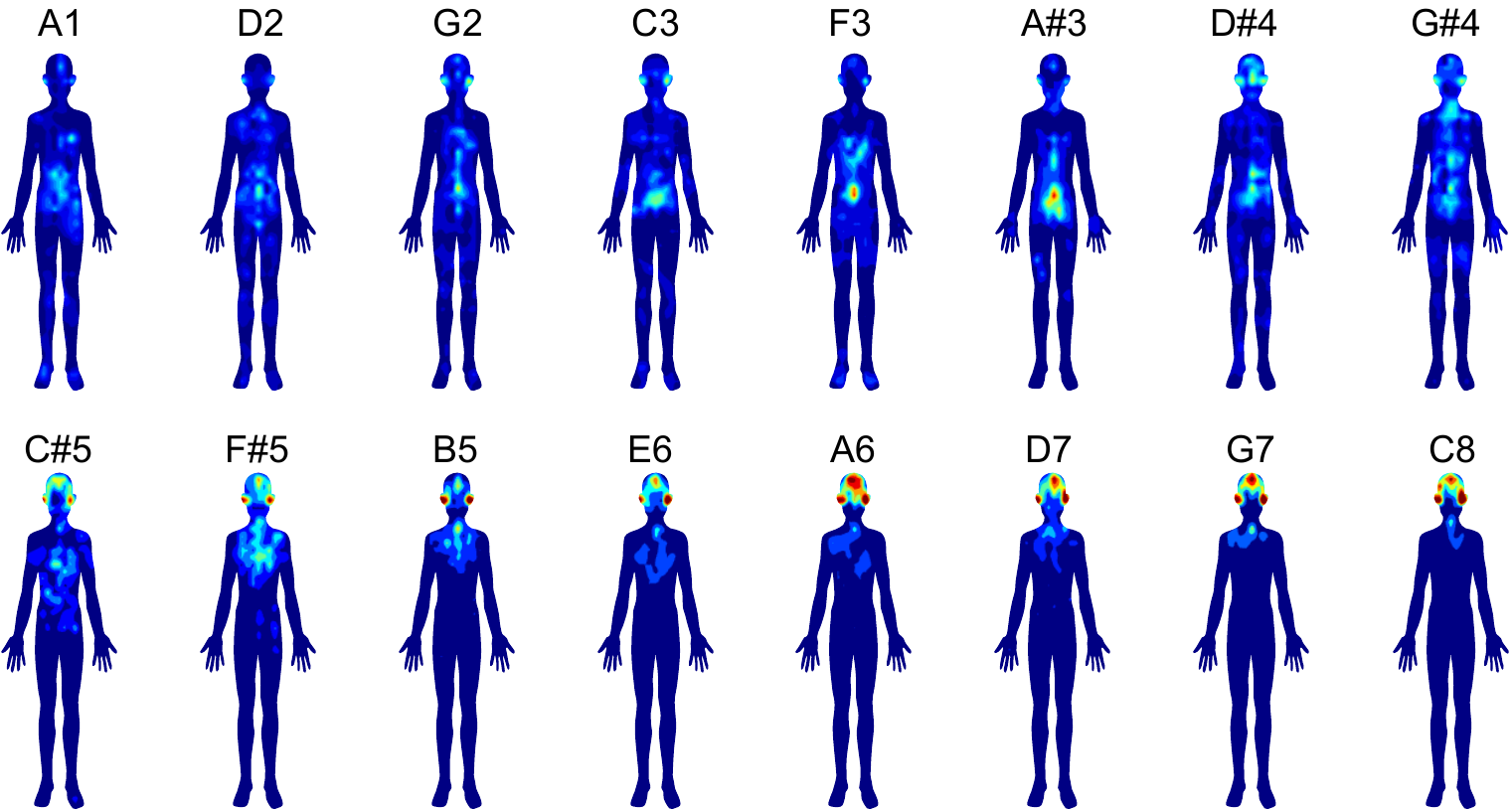

**Alexithymia Group**

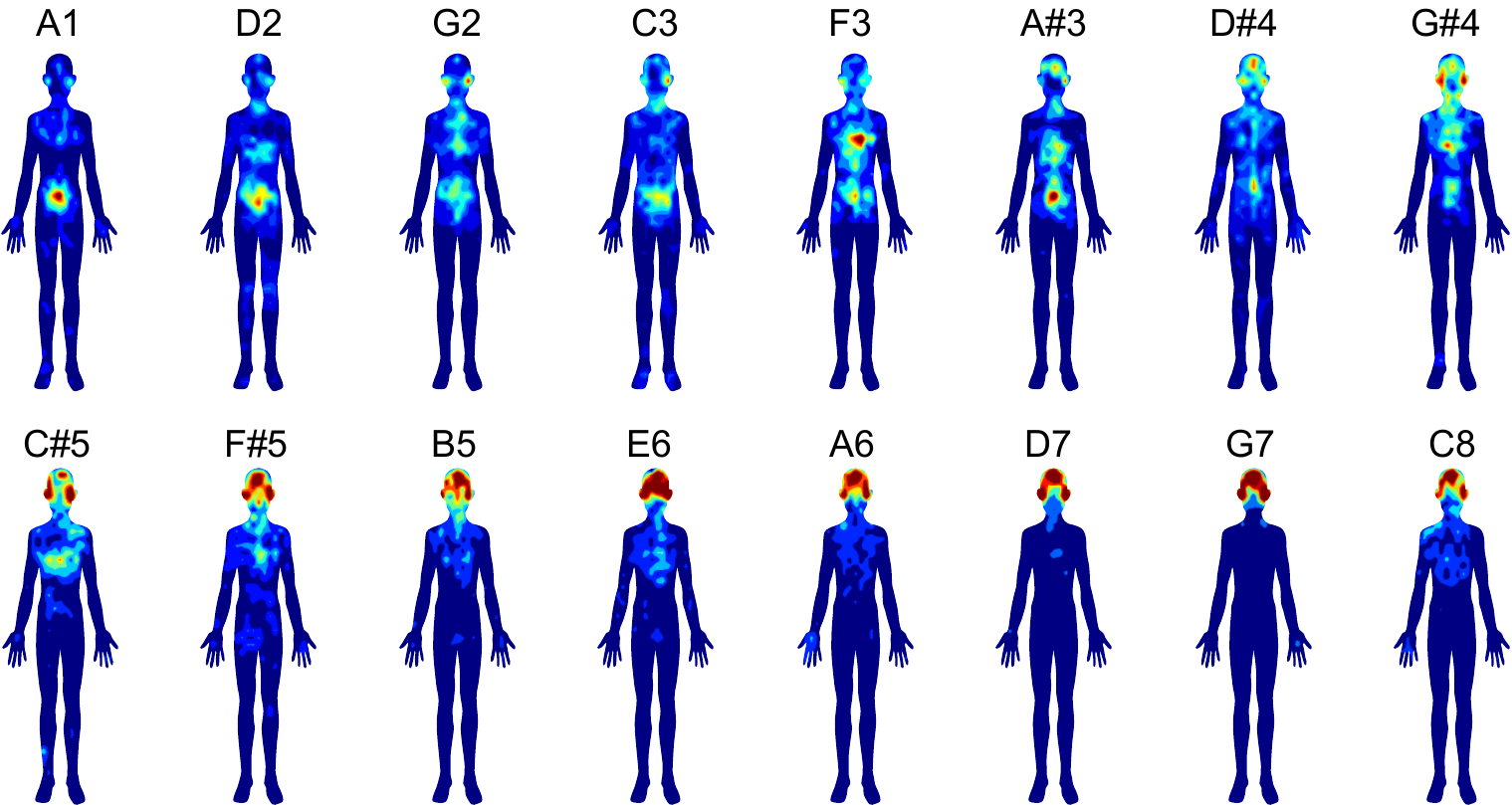

**Non alexithymia Group**

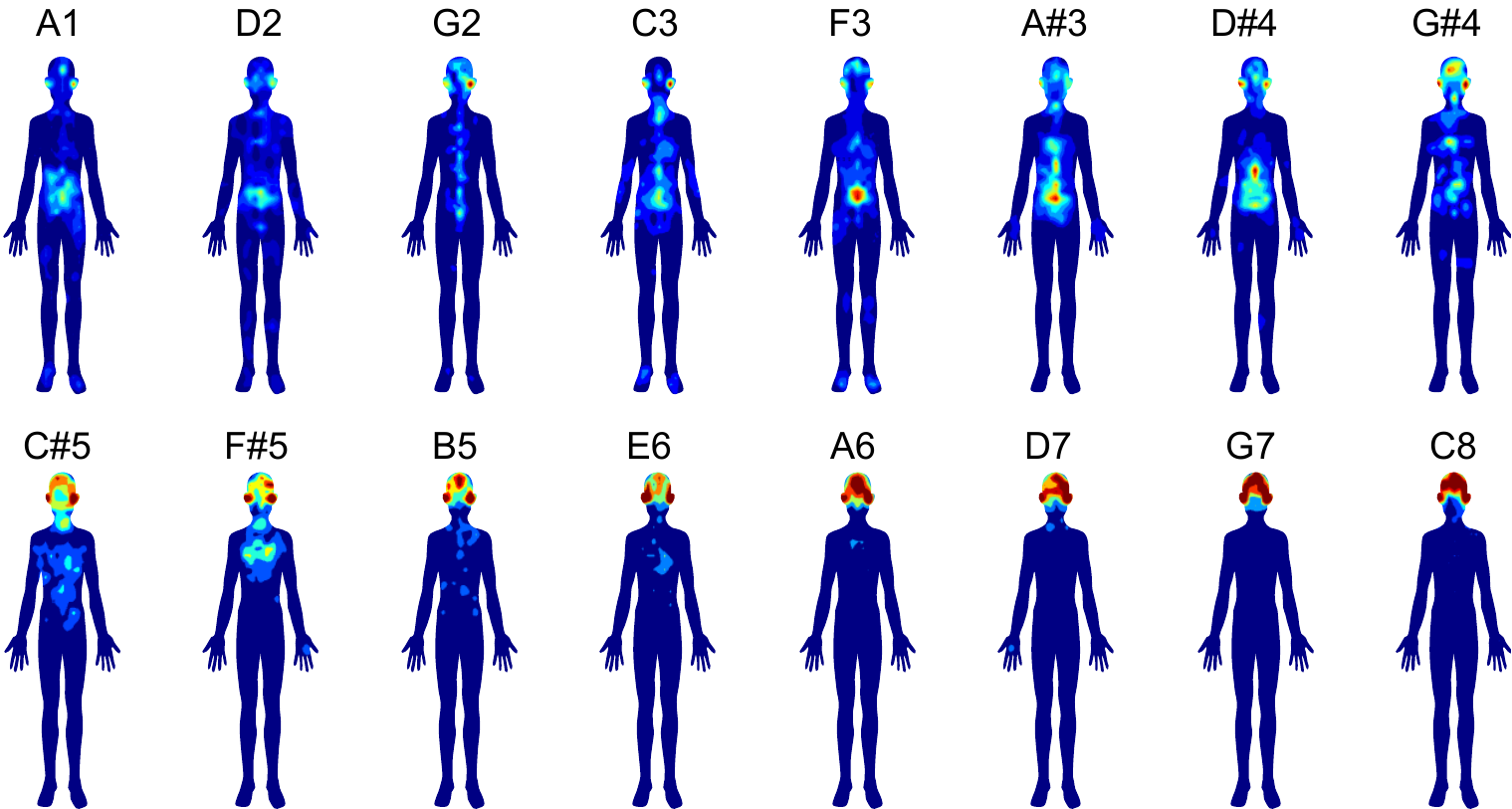

**Difficulty describing feelings (DDF) Group**

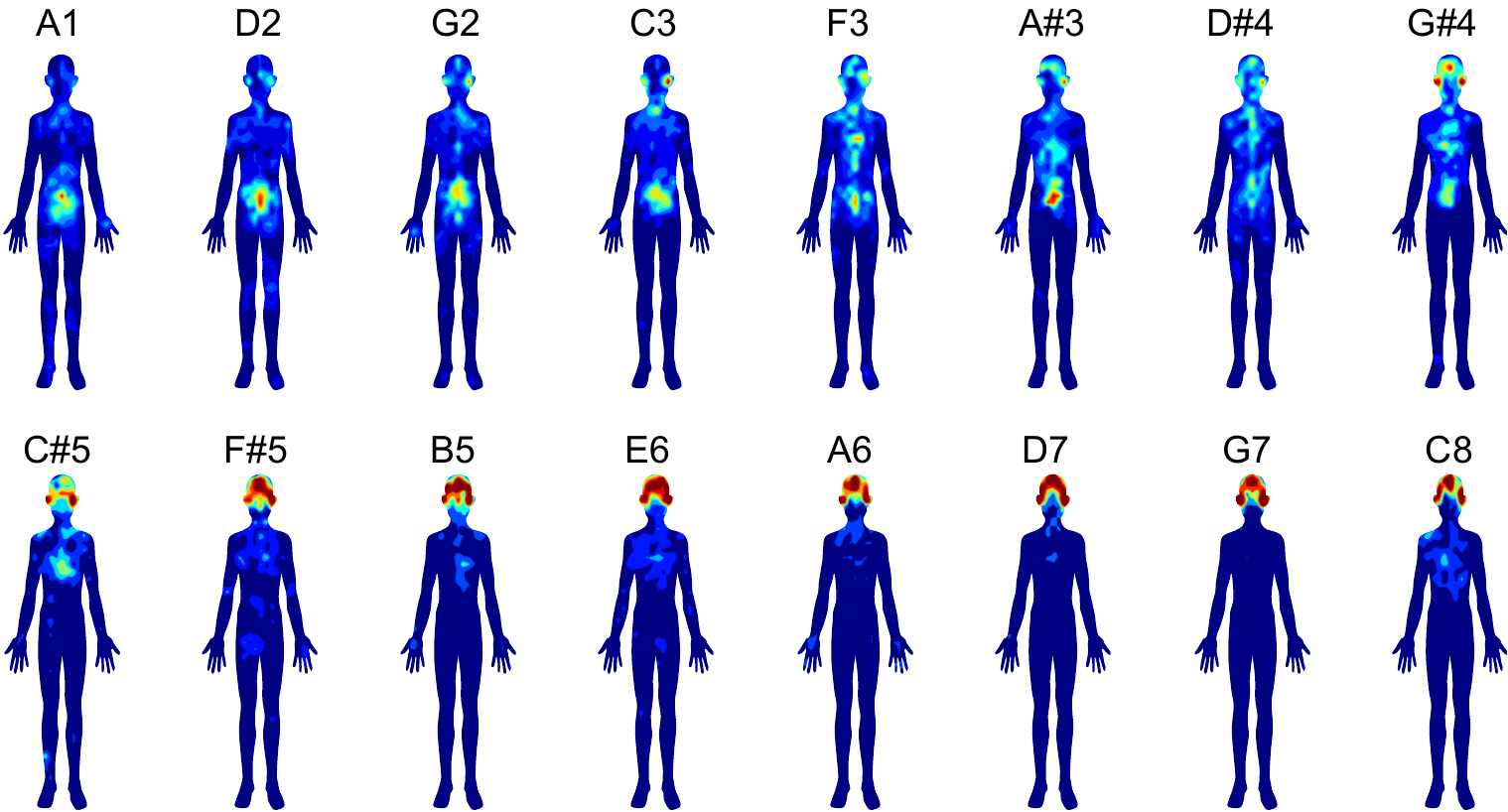

**Non difficulty describing feelings (Non-DDF) Group**

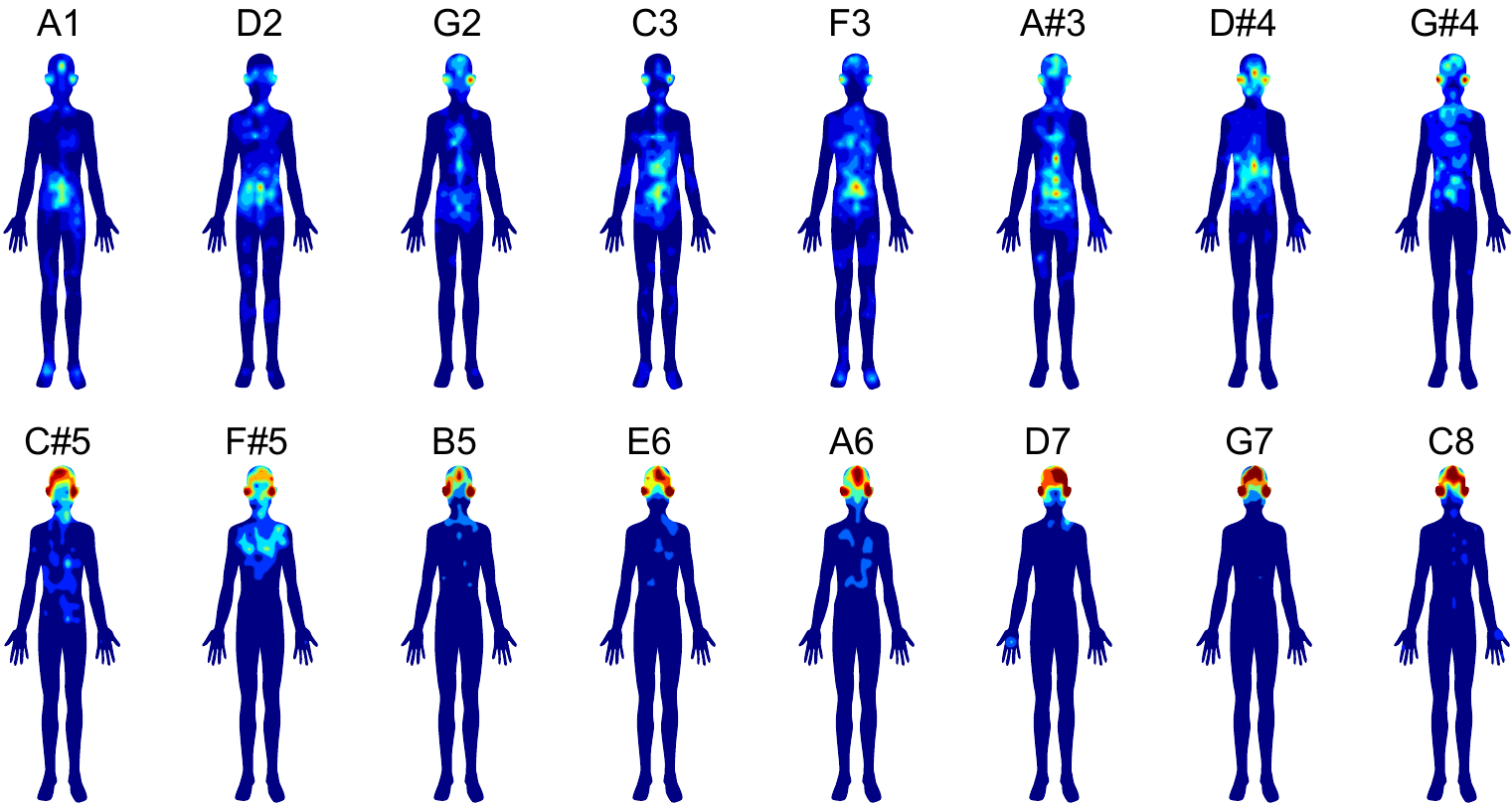

**Difficulty identifying feelings (DIF) Group**

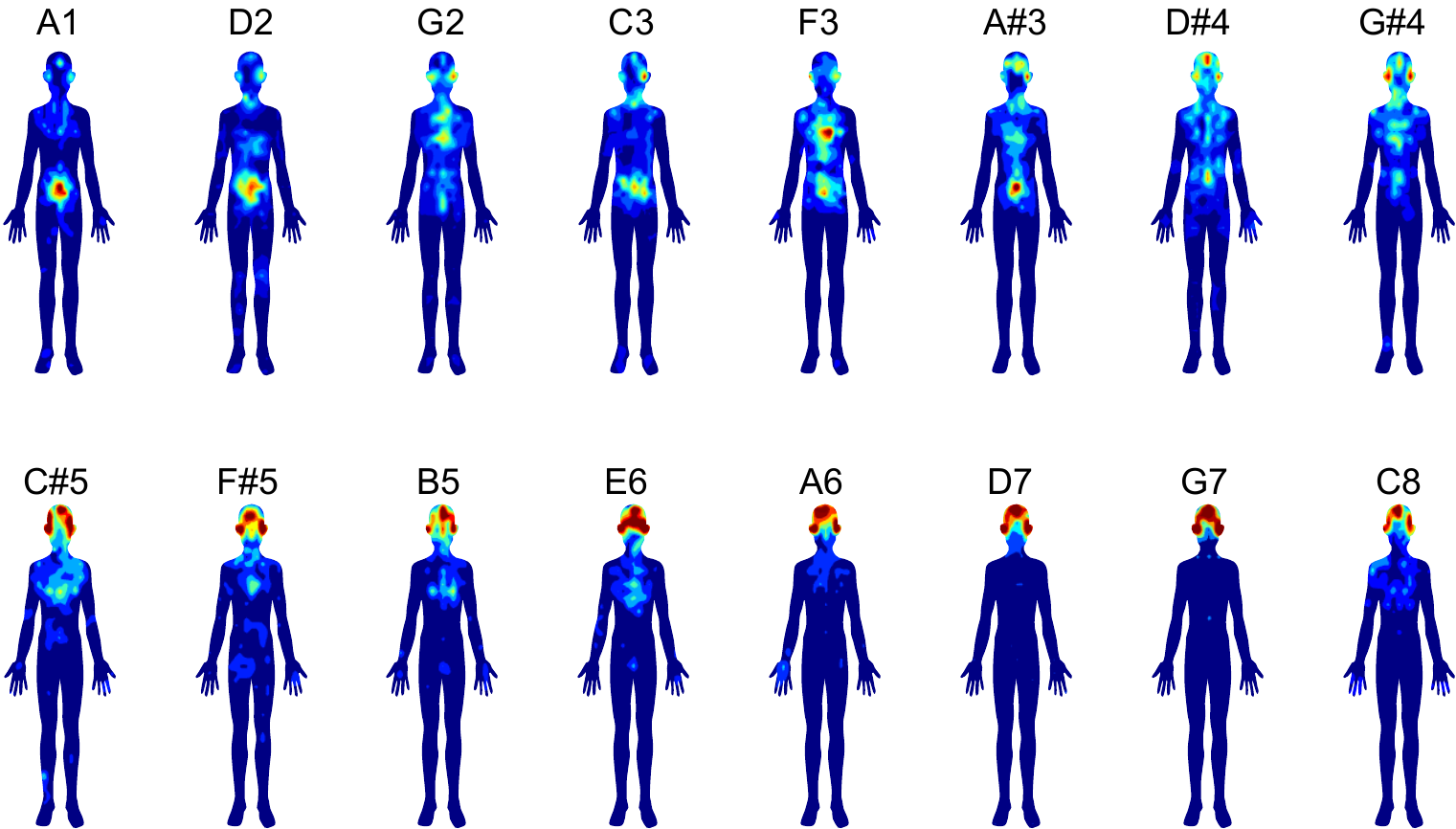

**Non difficulty identifying feelings (Non-DIF) Group**

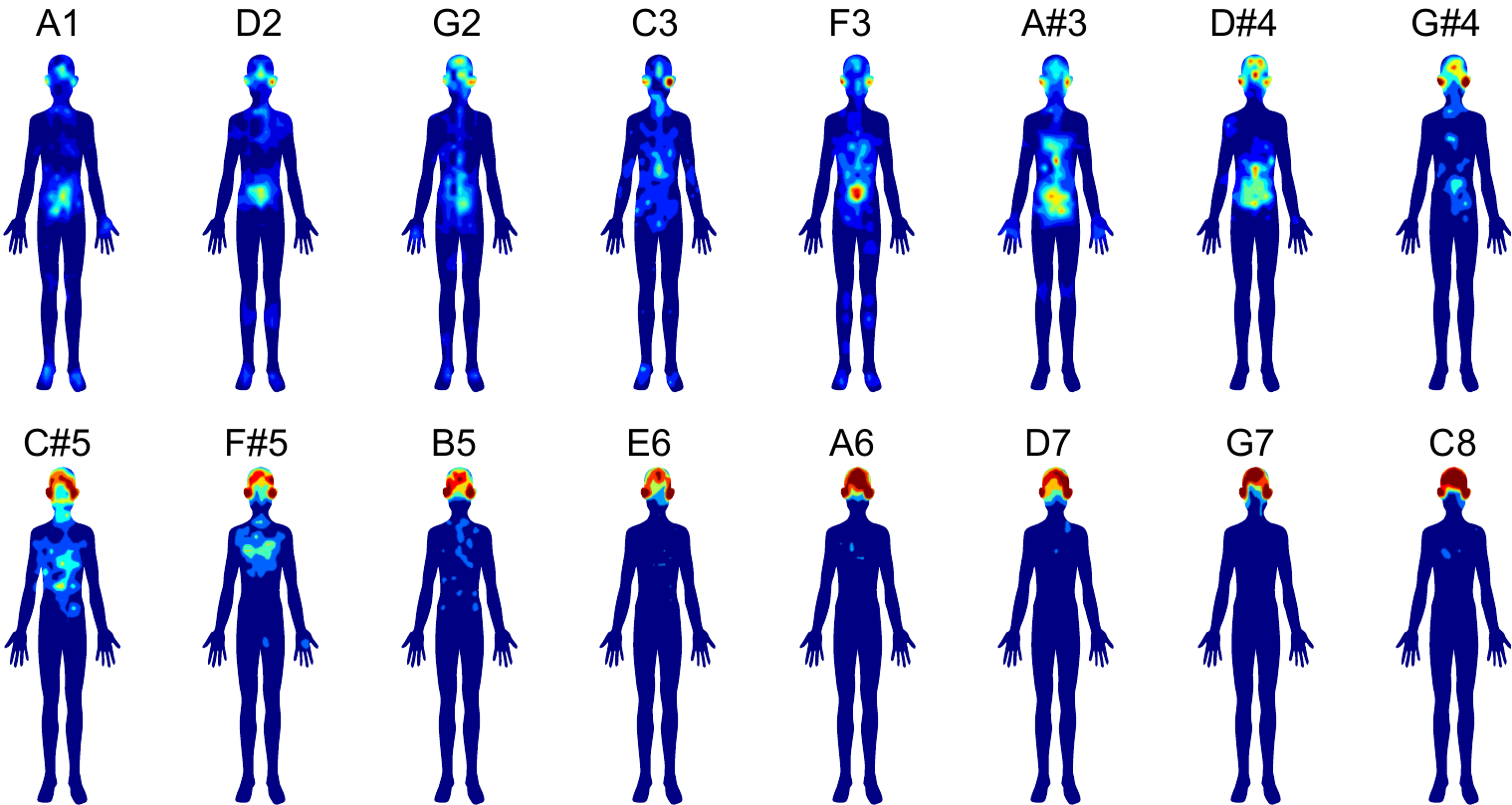

**Externally orientated thinking (EOT) Group**

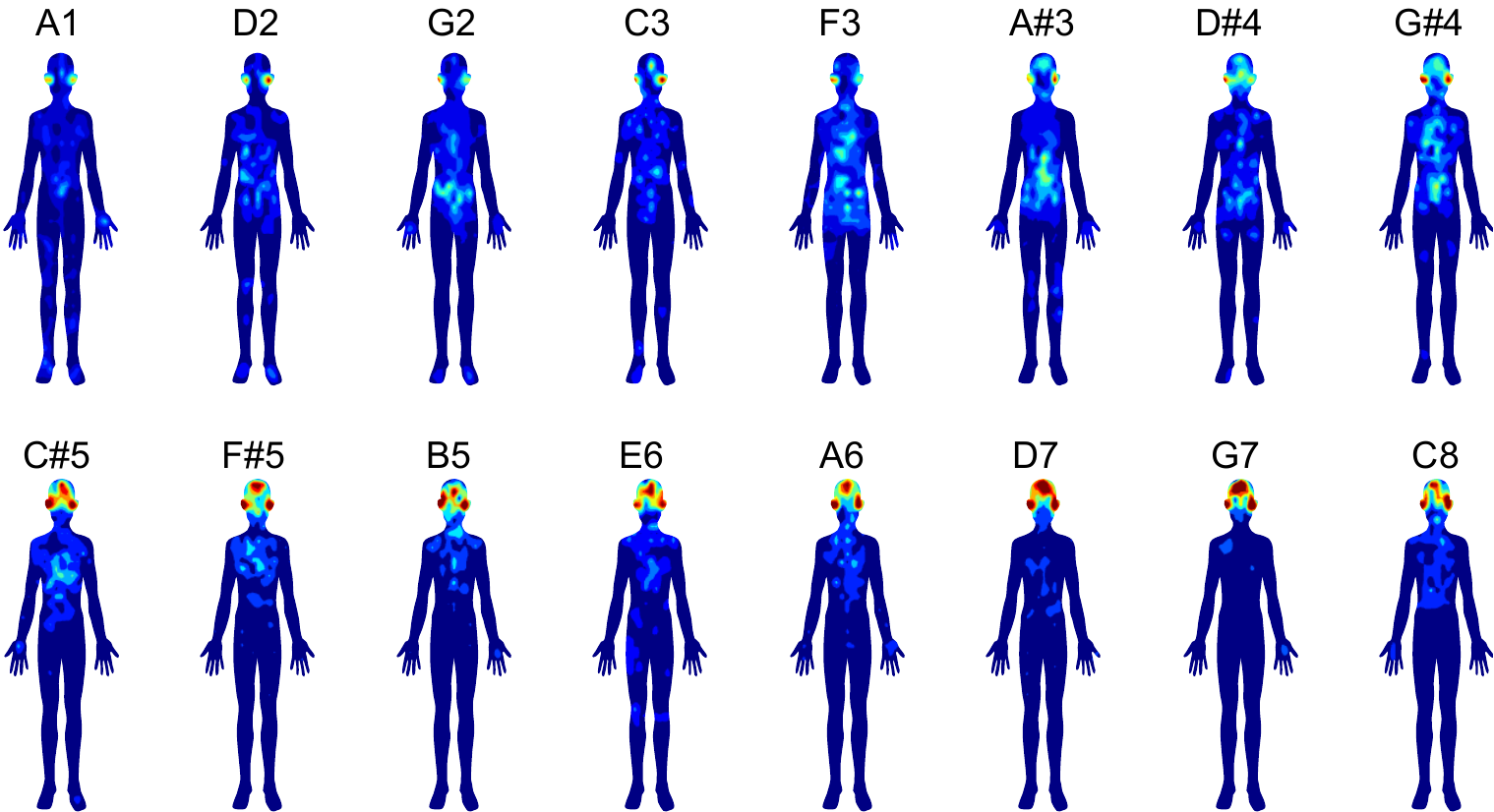

**Non externally orientated thinking (EOT) Group**

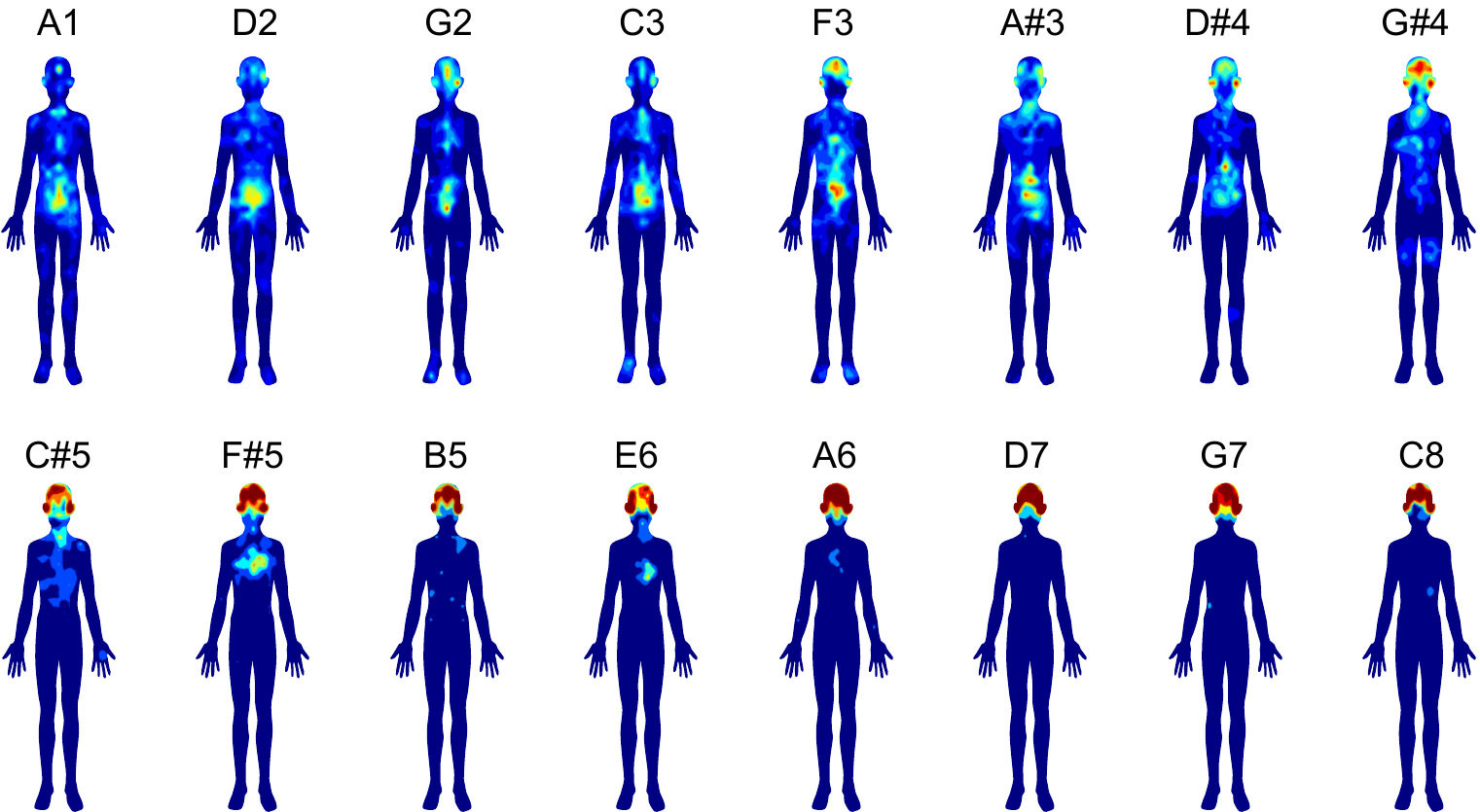

**Figure b. The result of pitch discrimination test**

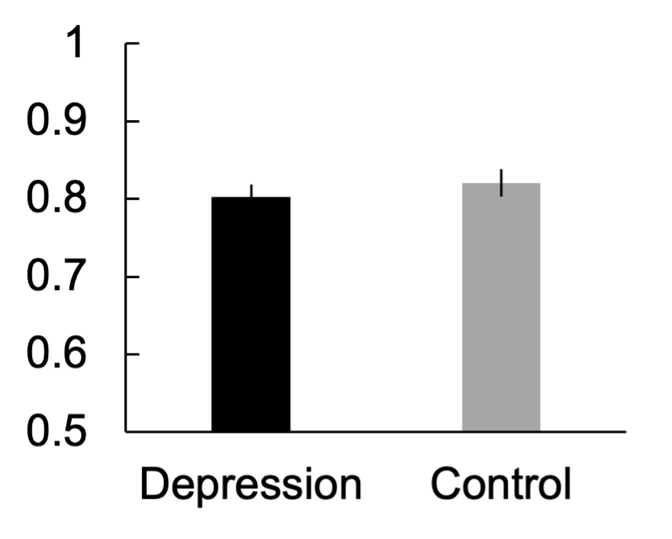

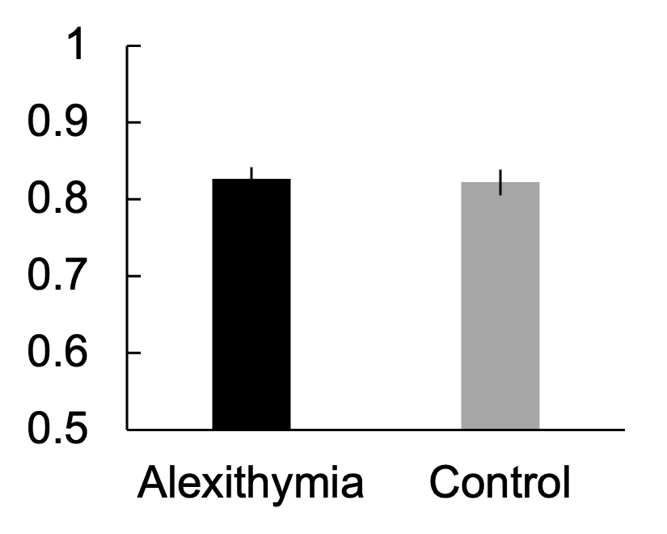
