## Supplementary material for "Body Maps of Sound Pitch and its individual differences in Alexithymia and Depression": S3 Appendix

**Figure. Valence and arousal for tone in each group.**

**Grand average of all participants**

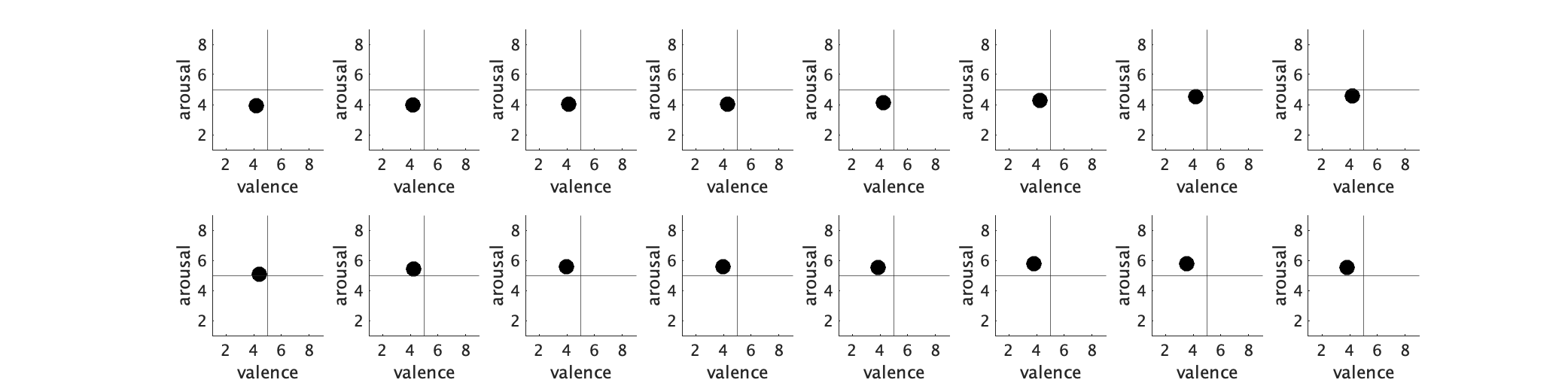

**Depression Group**

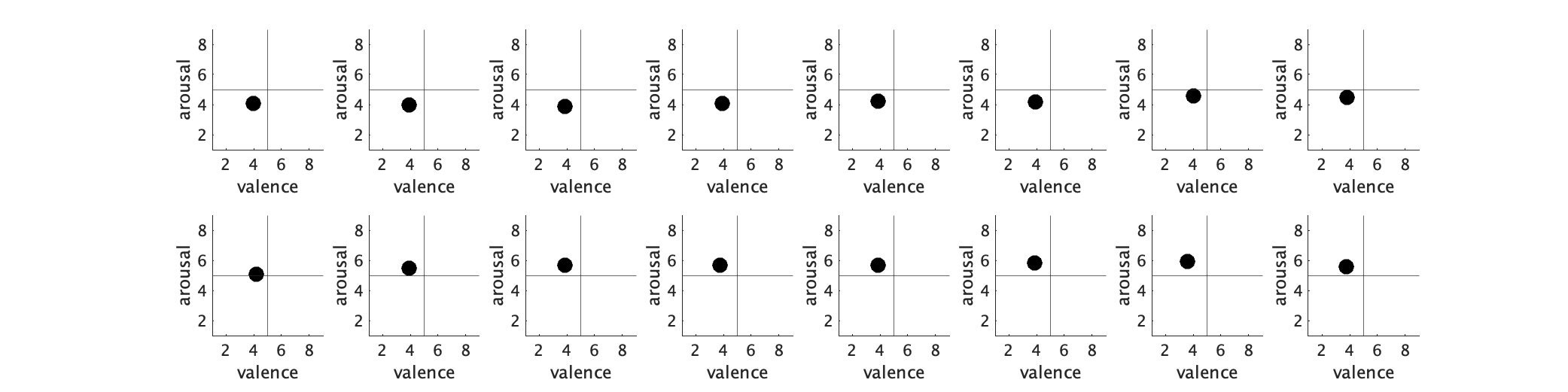

**Non depression Group**

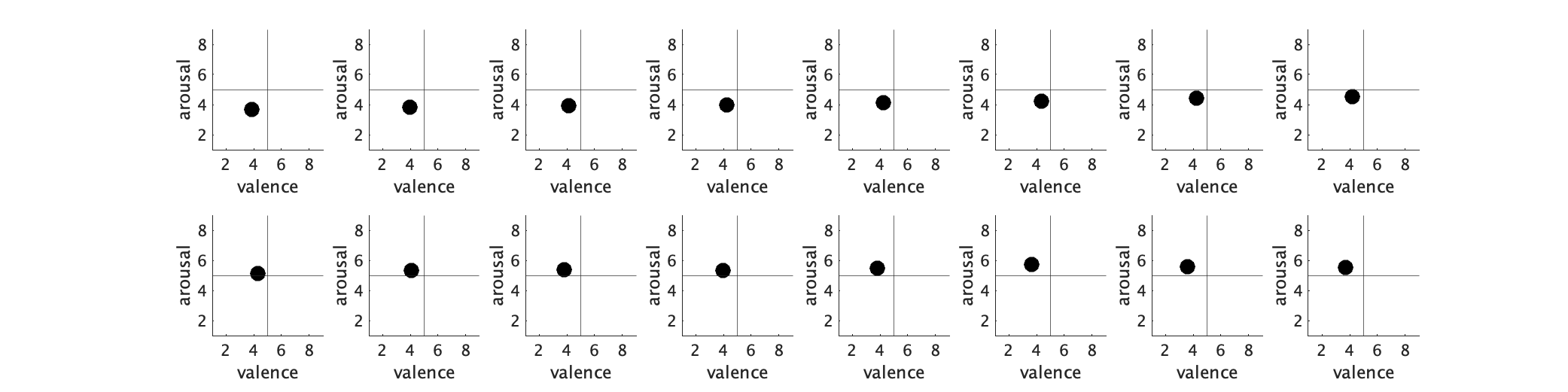

**Alexithymia Group**

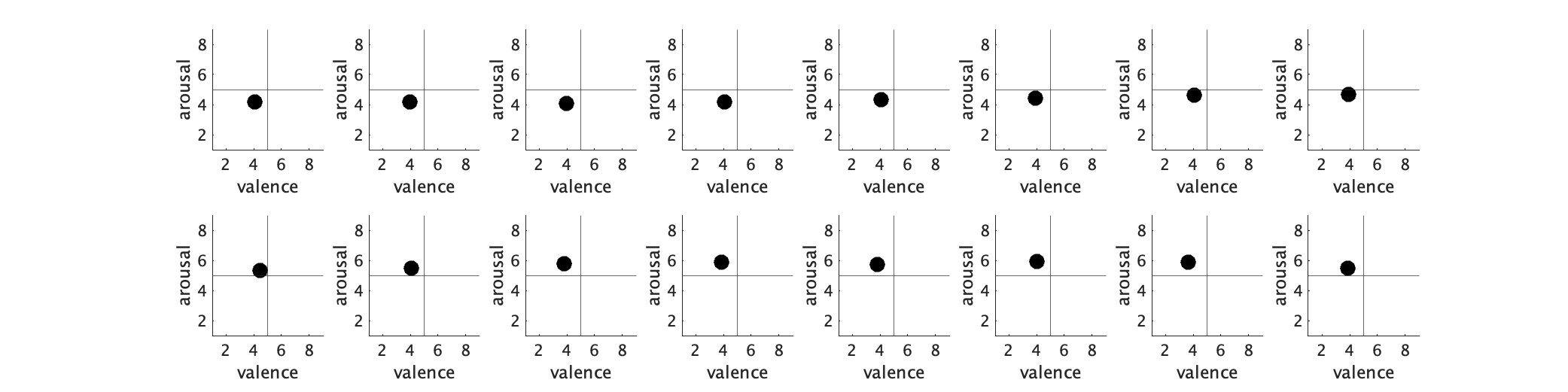

**Non alexithymia Group**

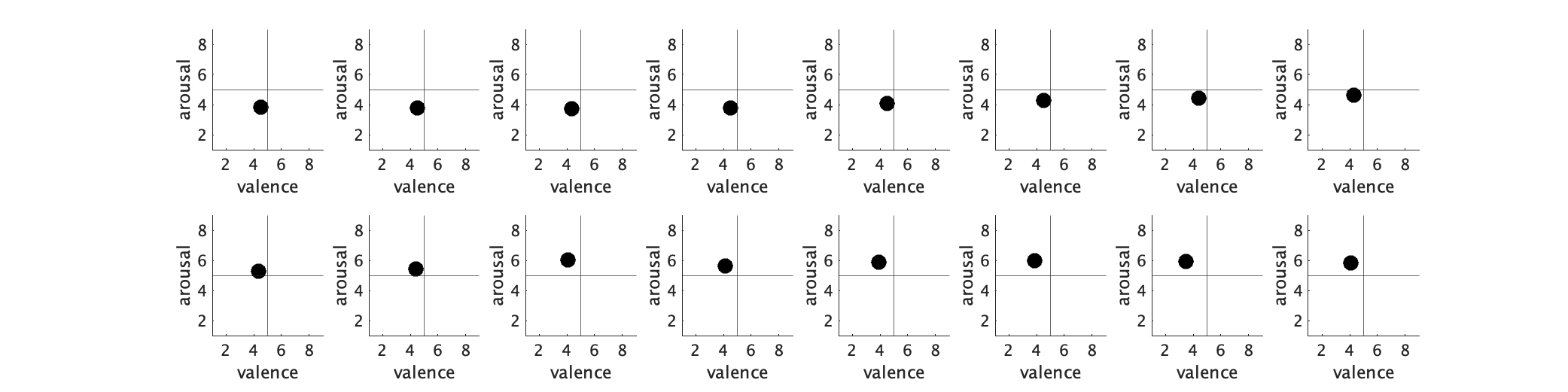

**Difficulty describing feelings (DDF) Group**

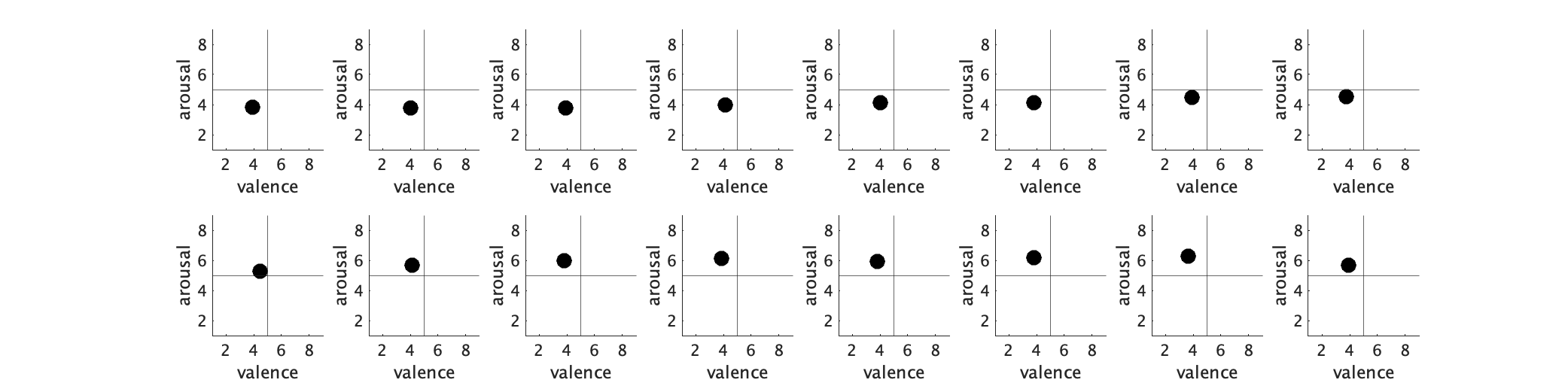

**Non difficulty describing feelings (Non-DDF) Group**

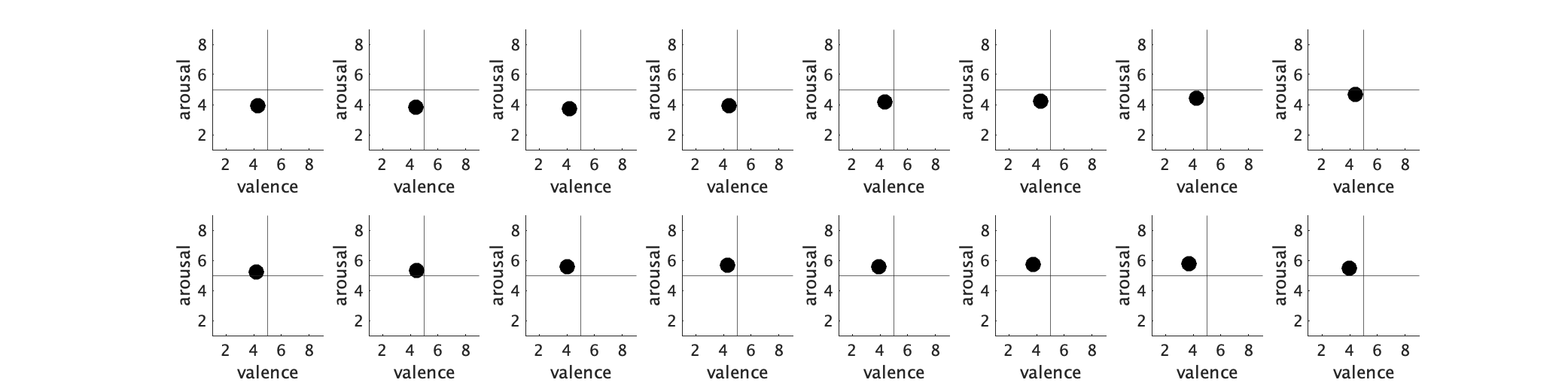

**Difficulty identifying feelings (DIF) Group**

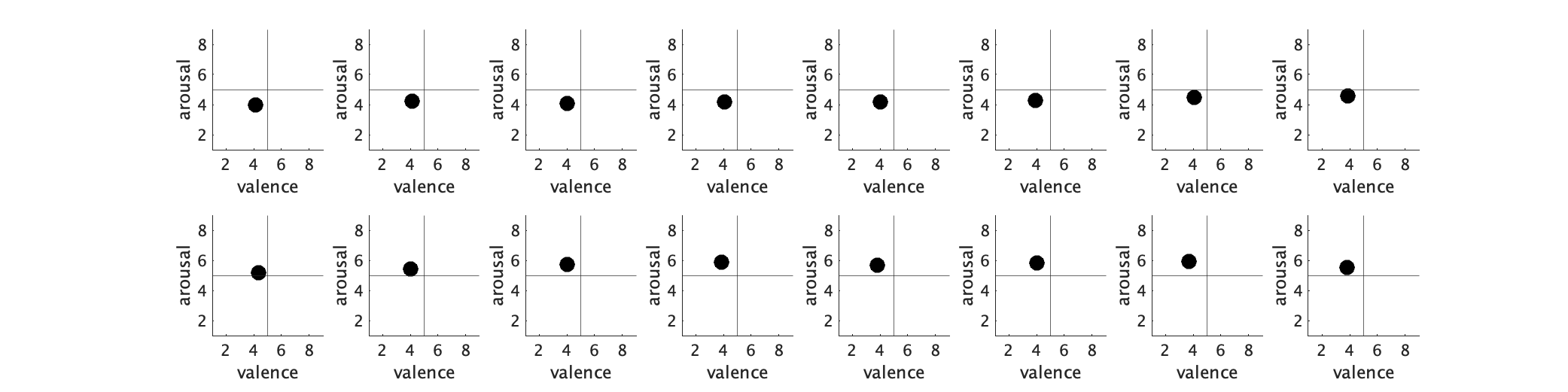

**Non difficulty identifying feelings (Non-DIF) Group**

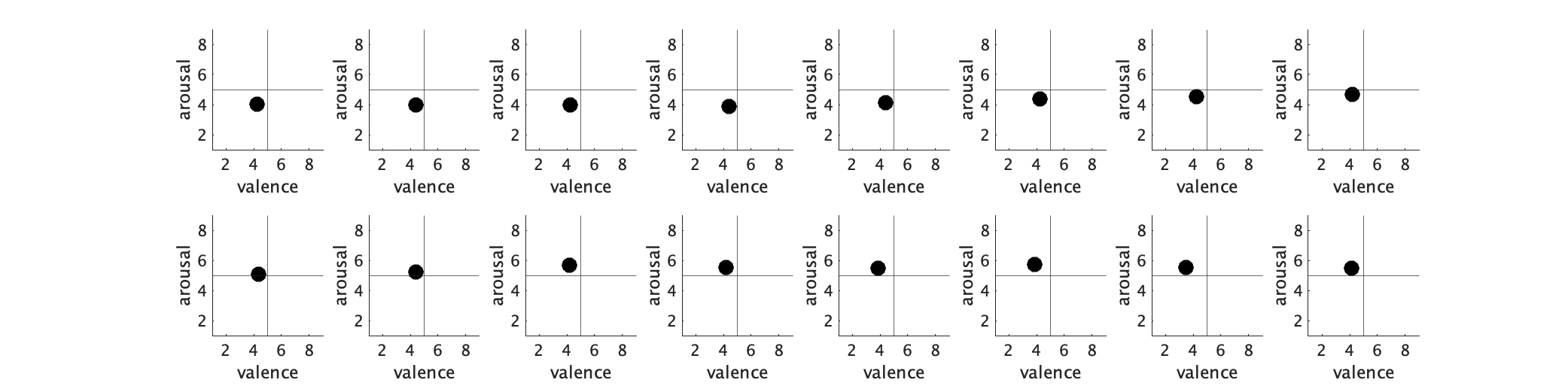

**Externally orientated thinking (EOT) Group**

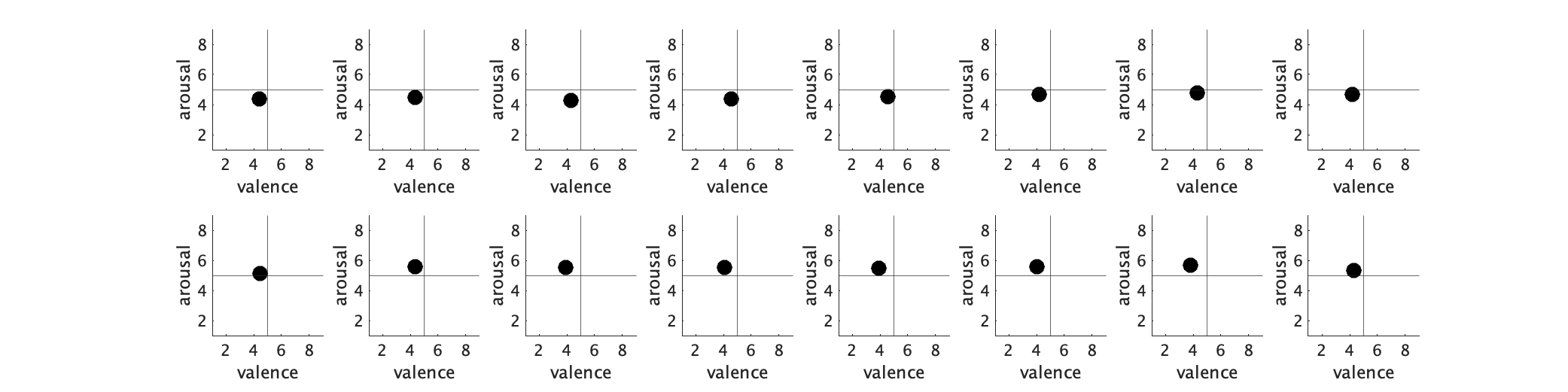

**Non externally orientated thinking (EOT) Group**

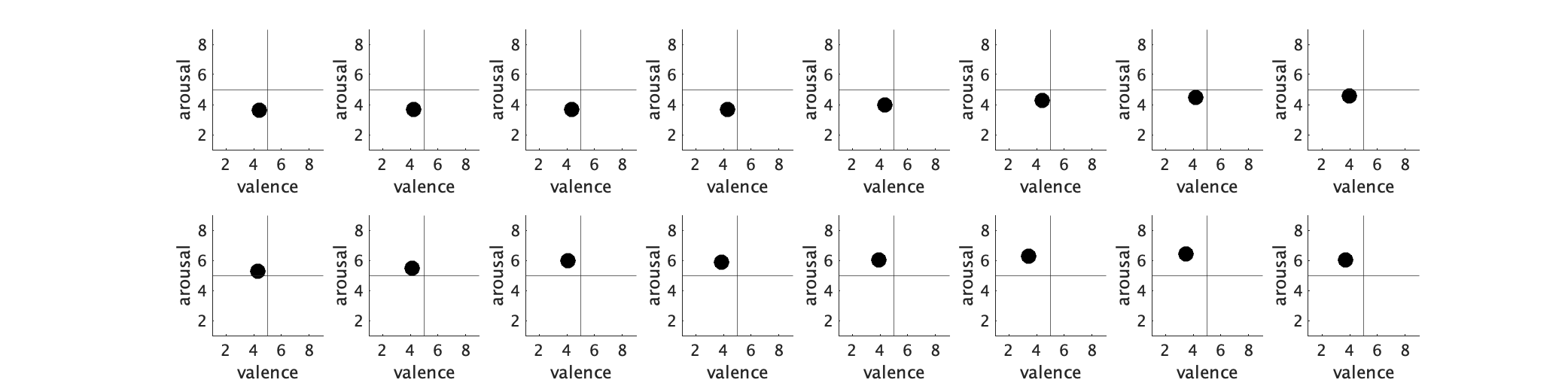
