## Supplementary material for "Body Maps of Sound Pitch and its individual differences in Alexithymia and Depression": S4 Appendix

**Table. 33 categories of emotion**

| 1 | Admiration |
| --- | --- |
| 2 | Adoration |
| 3 | Aesthetic appreciation |
| 4 | Amusement |
| 5 | Anger |
| 6 | Anxiety |
| 7 | Awe |
| 8 | Awkwardness |
| 9 | Boredom |
| 10 | Calmness |
| 11 | Confusion |
| 12 | Contempt |
| 13 | Craving |
| 14 | Disappointment |
| 15 | Disgust |
| 16 | Empathy |
| 17 | Entrancement |
| 18 | Envy |
| 19 | Excitement |
| 20 | Fear/Horror |
| 21 | Guilt |
| 22 | Interest |
| 23 | Joy |
| 24 | Nostalgia |
| 25 | Pride |
| 26 | Relief |
| 27 | Romance |
| 28 | Sadness |
| 29 | Satisfaction |
| 30 | Sexual desire |
| 31 | Surprise |
| 32 | Sympathy |
| 33 | Triumph |

**Figure a. Categorical judgement of 33 emotions for tone in each group.**

**Grand average of all participants**

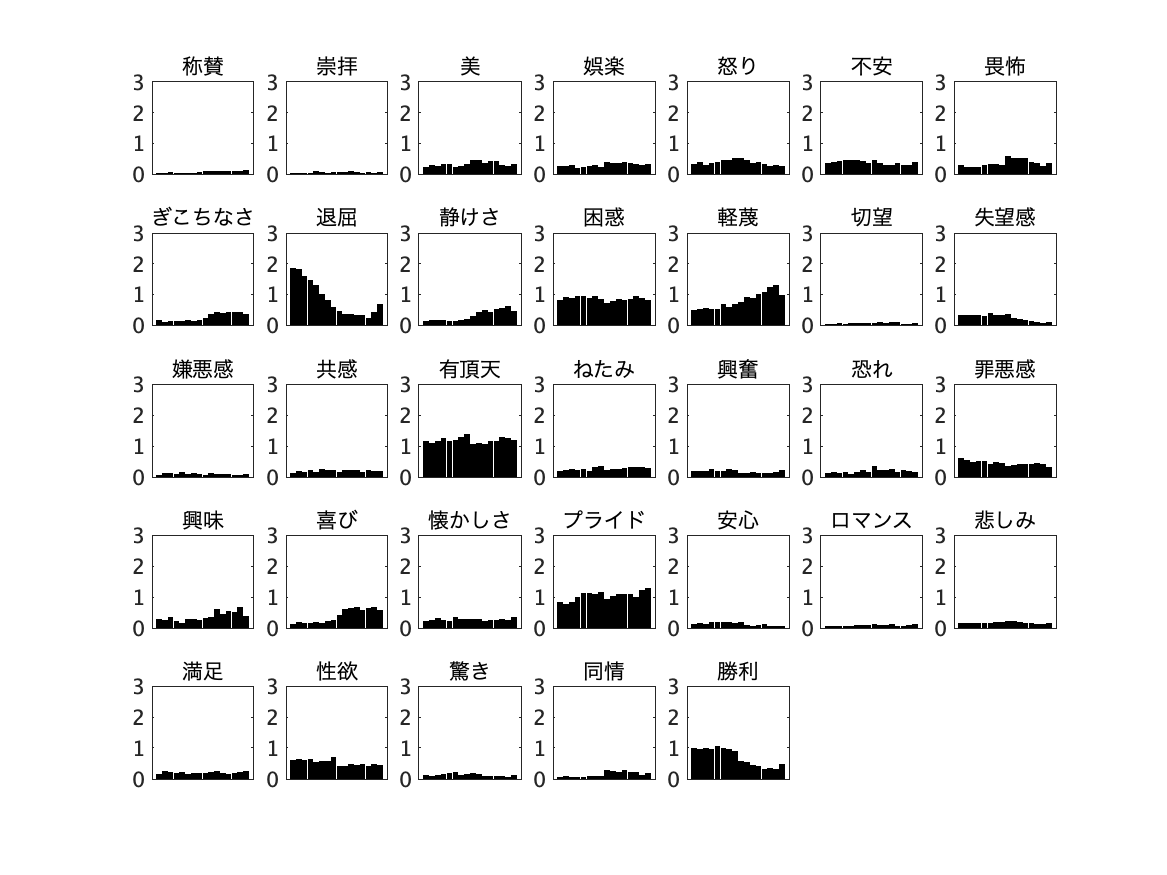

Admiration	Adoration	Aesthetic	Amusement	Anger	Anxiety	Awe

Awkwardness	Boredom	Calmness	Confusion	Contempt	Craving	Disappointment

Disgust	Empathy Entrancement	Envy	Excitement	Fear/Horror	Guilt

Interest	Joy	Nostalgia	Pride	Relief	Romance	Sadness

Satisfaction Sexual desire	Surprise	Sympathy	Triumph

**Depression Group**

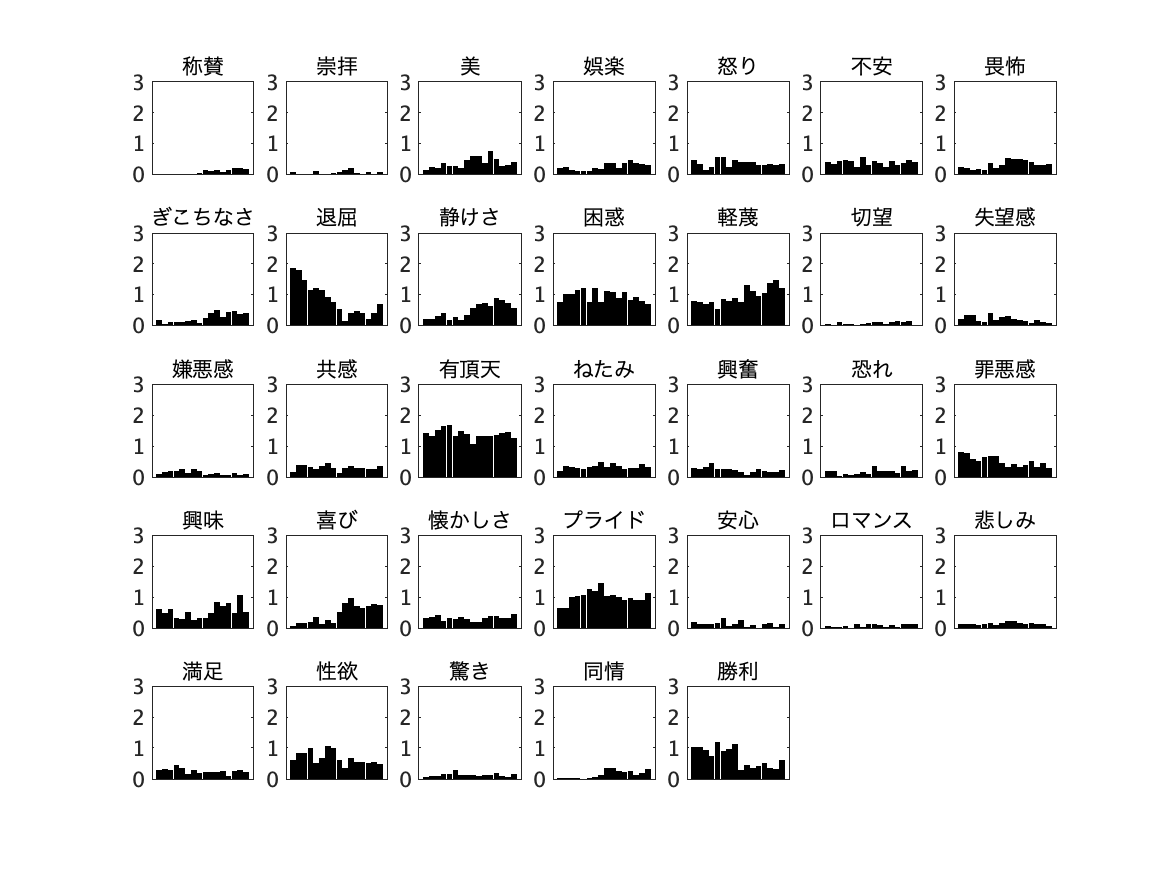

Admiration	Adoration	Aesthetic	Amusement	Anger	Anxiety	Awe

Satisfaction Sexual desire	Surprise	Sympathy	Triumph

Awkwardness	Boredom	Calmness	Confusion	Contempt	Craving	Disappointment

Disgust	Empathy Entrancement	Envy	Excitement	Fear/Horror	Guilt

Interest	Joy	Nostalgia	Pride	Relief	Romance	Sadness

**Non depression Group**

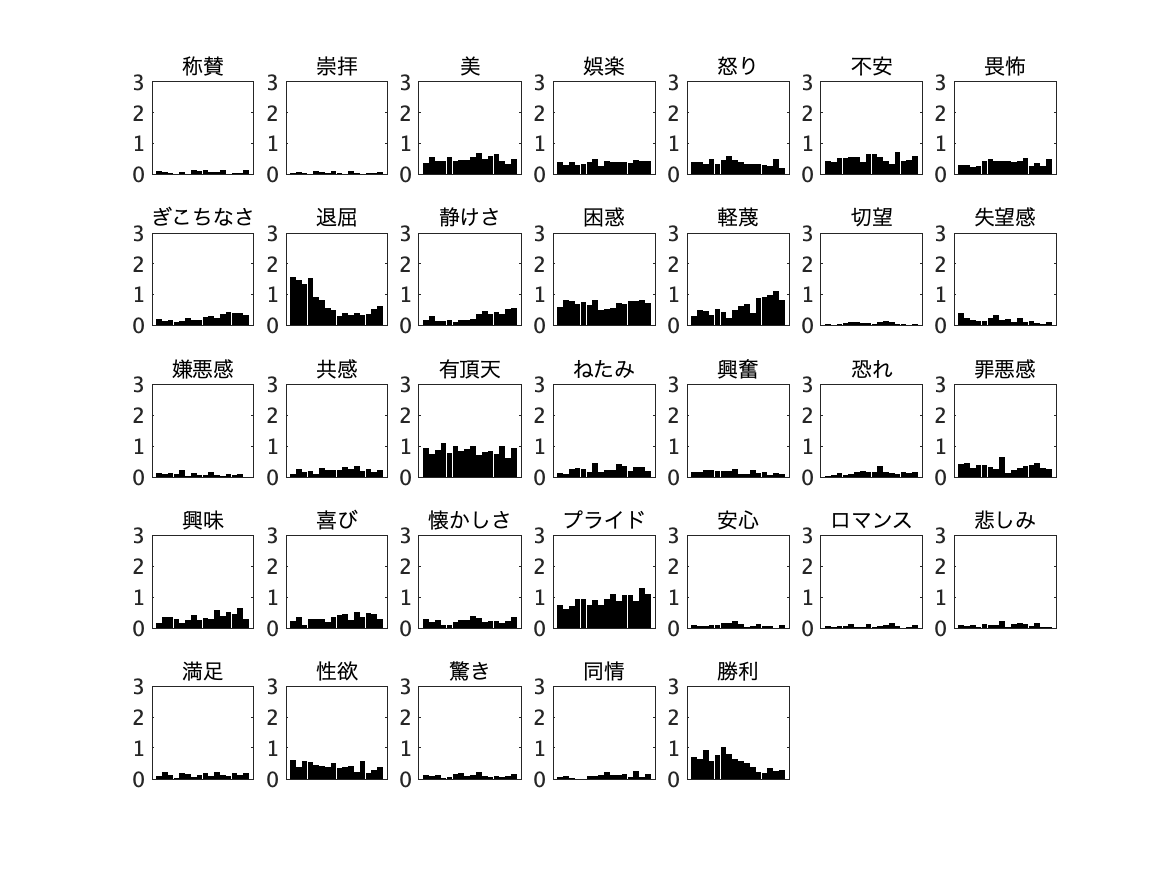

Admiration	Adoration	Aesthetic	Amusement	Anger	Anxiety	Awe

Awkwardness	Boredom	Calmness	Confusion	Contempt	Craving	Disappointment

Disgust	Empathy Entrancement	Envy	Excitement	Fear/Horror	Guilt

Interest	Joy	Nostalgia	Pride	Relief	Romance	Sadness

Satisfaction Sexual desire	Surprise	Sympathy	Triumph

**Alexithymia Group**

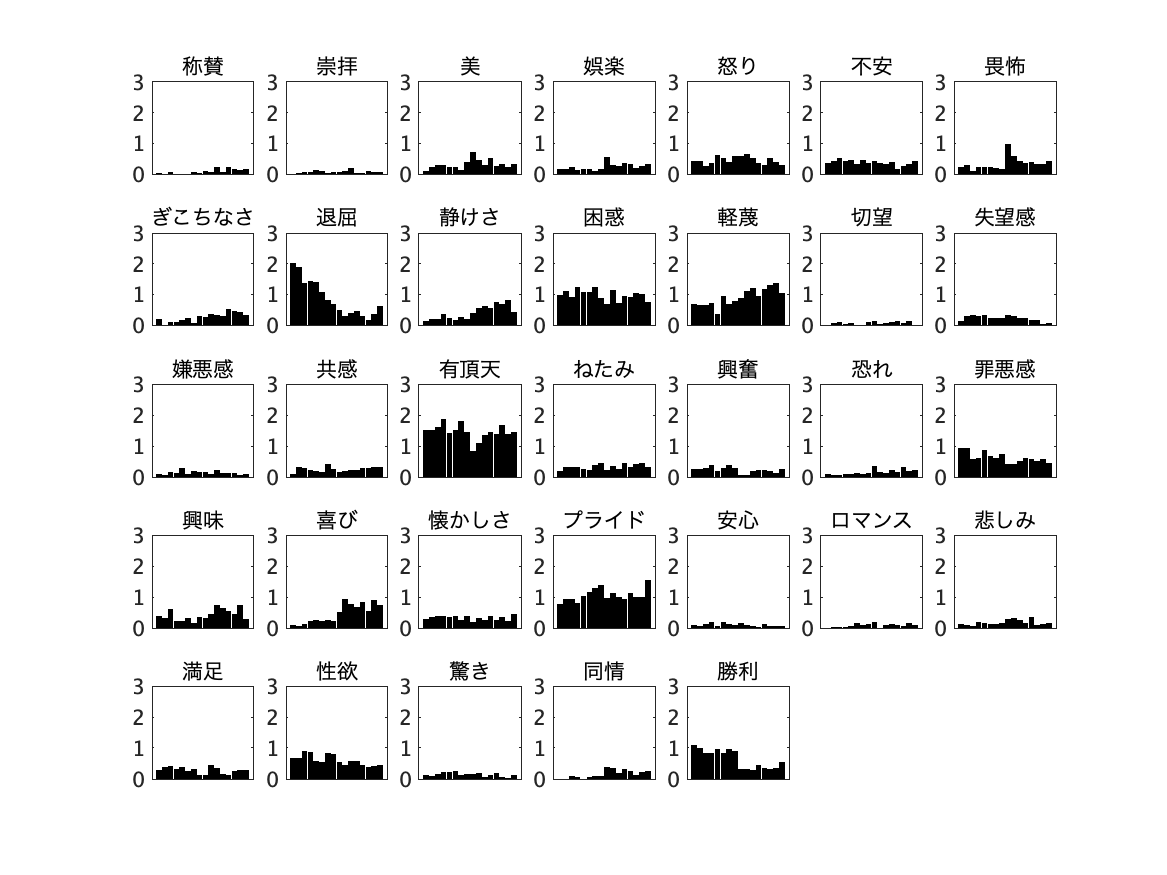

Admiration	Adoration	Aesthetic	Amusement	Anger	Anxiety	Awe

Awkwardness	Boredom	Calmness	Confusion	Contempt	Craving	Disappointment

Disgust	Empathy Entrancement	Envy	Excitement	Fear/Horror	Guilt

Interest	Joy	Nostalgia	Pride	Relief	Romance	Sadness

Satisfaction Sexual desire	Surprise	Sympathy	Triumph

**Non alexithymia Group**

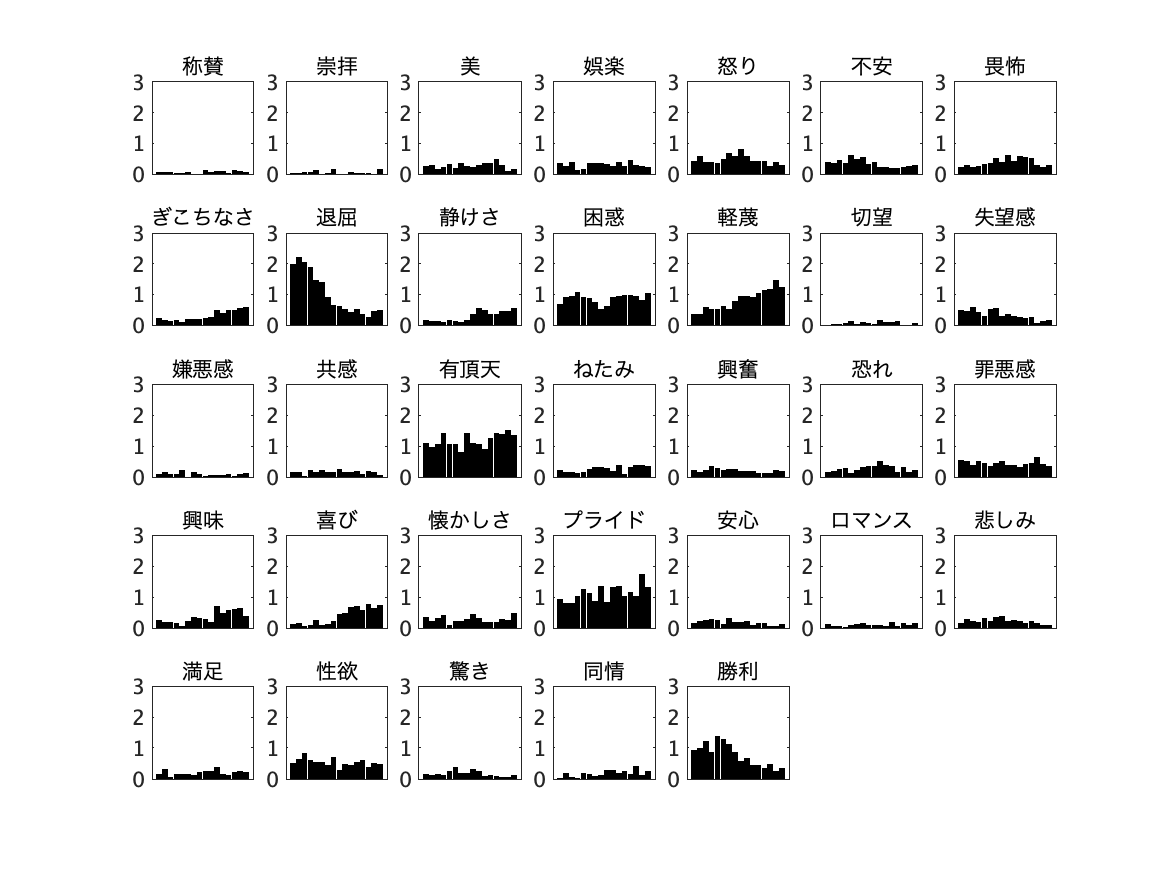

Admiration	Adoration	Aesthetic	Amusement	Anger	Anxiety	Awe

Satisfaction Sexual desire	Surprise	Sympathy	Triumph

Interest	Joy	Nostalgia	Pride	Relief	Romance	Sadness

Disgust	Empathy Entrancement	Envy	Excitement	Fear/Horror	Guilt

Awkwardness	Boredom	Calmness	Confusion	Contempt	Craving	Disappointment

**Difficulty describing feelings (DDF) Group**

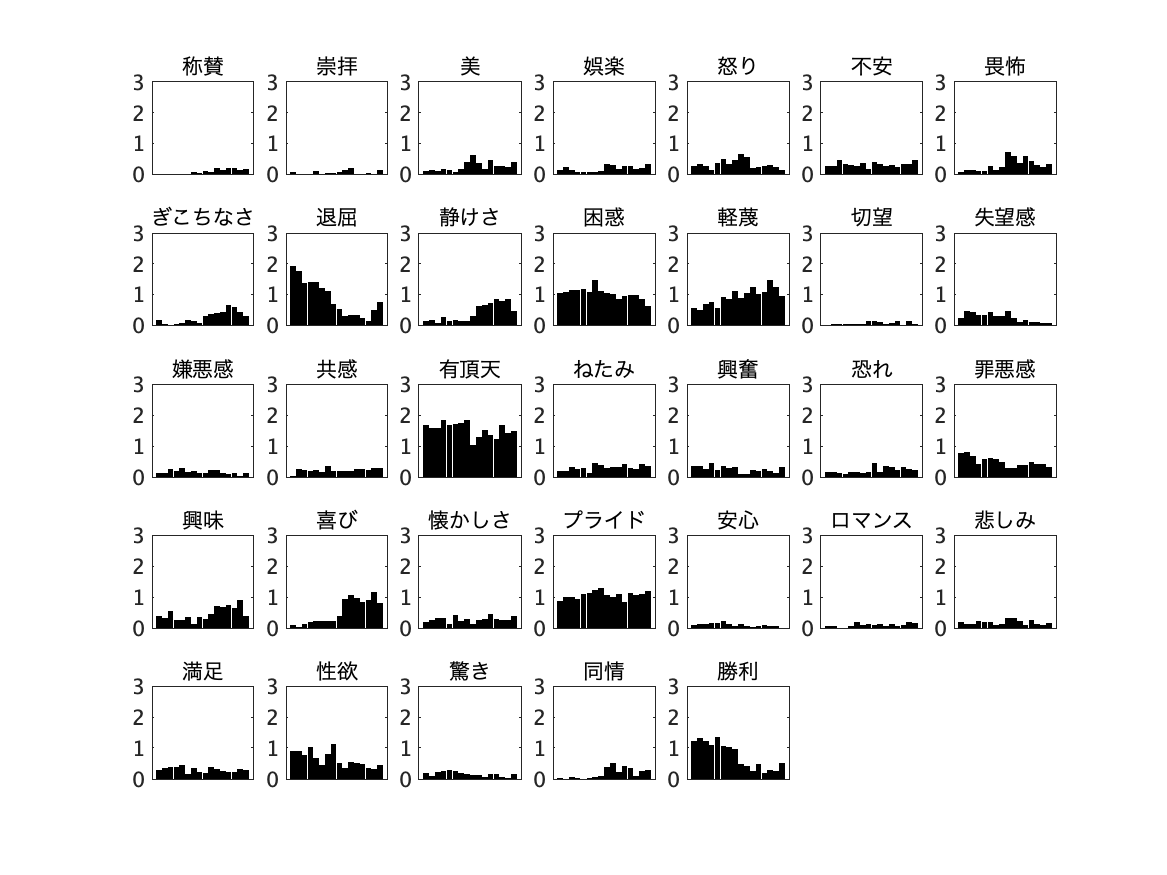

Admiration	Adoration	Aesthetic	Amusement	Anger	Anxiety	Awe

Awkwardness	Boredom	Calmness	Confusion	Contempt	Craving	Disappointment

Disgust	Empathy Entrancement	Envy	Excitement	Fear/Horror	Guilt

Interest	Joy	Nostalgia	Pride	Relief	Romance	Sadness

Satisfaction Sexual desire	Surprise	Sympathy	Triumph

**Non difficulty describing feelings (Non-DDF) Group**

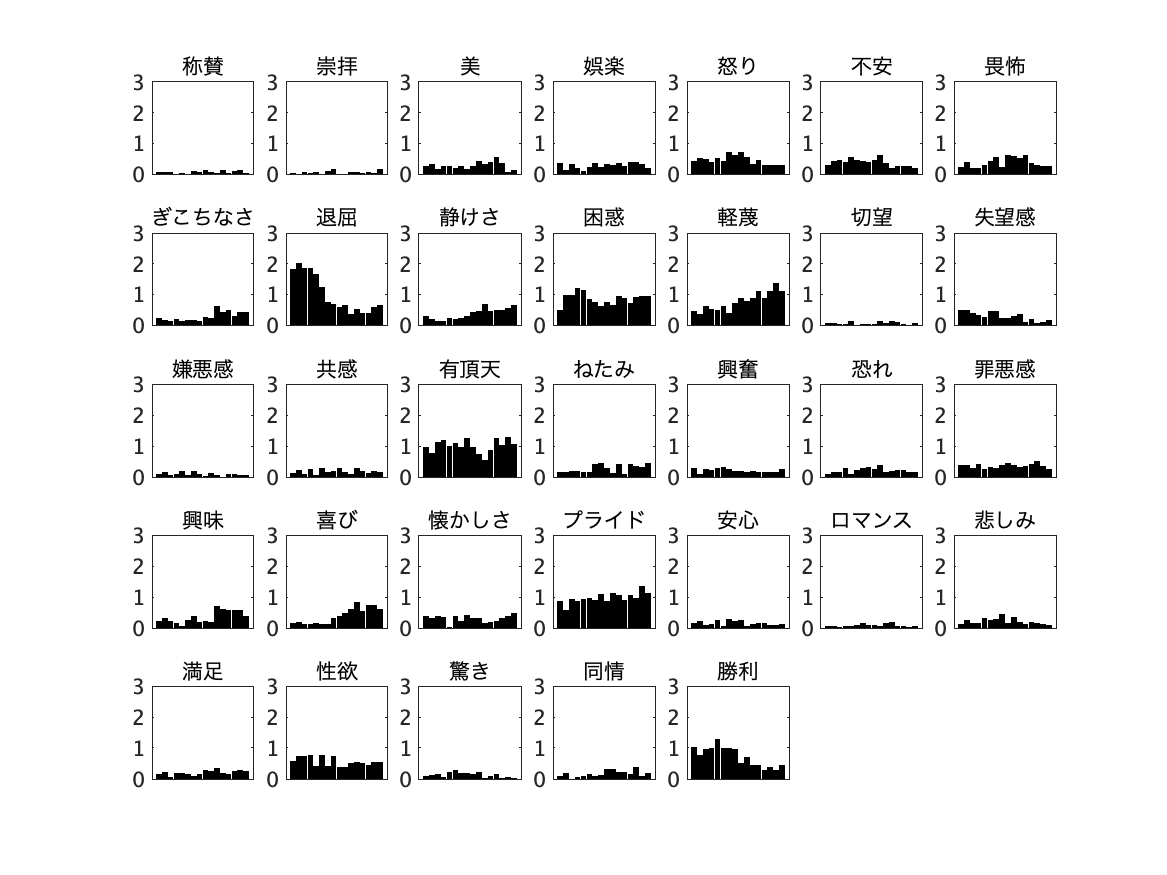

Admiration	Adoration	Aesthetic	Amusement	Anger	Anxiety	Awe

Awkwardness	Boredom	Calmness	Confusion	Contempt	Craving	Disappointment

Disgust	Empathy Entrancement	Envy	Excitement	Fear/Horror	Guilt

Interest	Joy	Nostalgia	Pride	Relief	Romance	Sadness

Satisfaction Sexual desire	Surprise	Sympathy	Triumph

**Difficulty identifying feelings (DIF) Group**

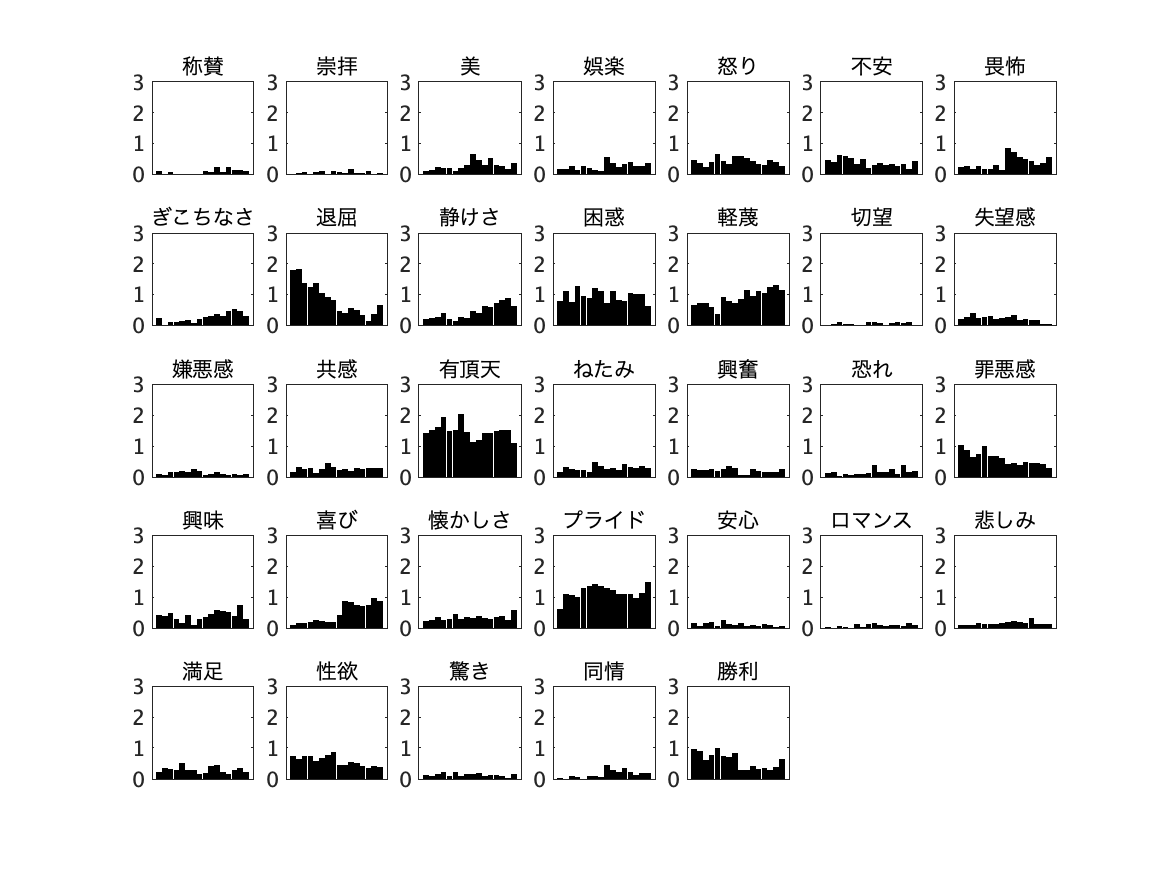

Admiration	Adoration	Aesthetic	Amusement	Anger	Anxiety	Awe

Awkwardness	Boredom	Calmness	Confusion	Contempt	Craving	Disappointment

Disgust	Empathy Entrancement	Envy	Excitement	Fear/Horror	Guilt

Interest	Joy	Nostalgia	Pride	Relief	Romance	Sadness

Satisfaction Sexual desire	Surprise	Sympathy	Triumph

**Non difficulty identifying feelings (Non-DIF) Group**

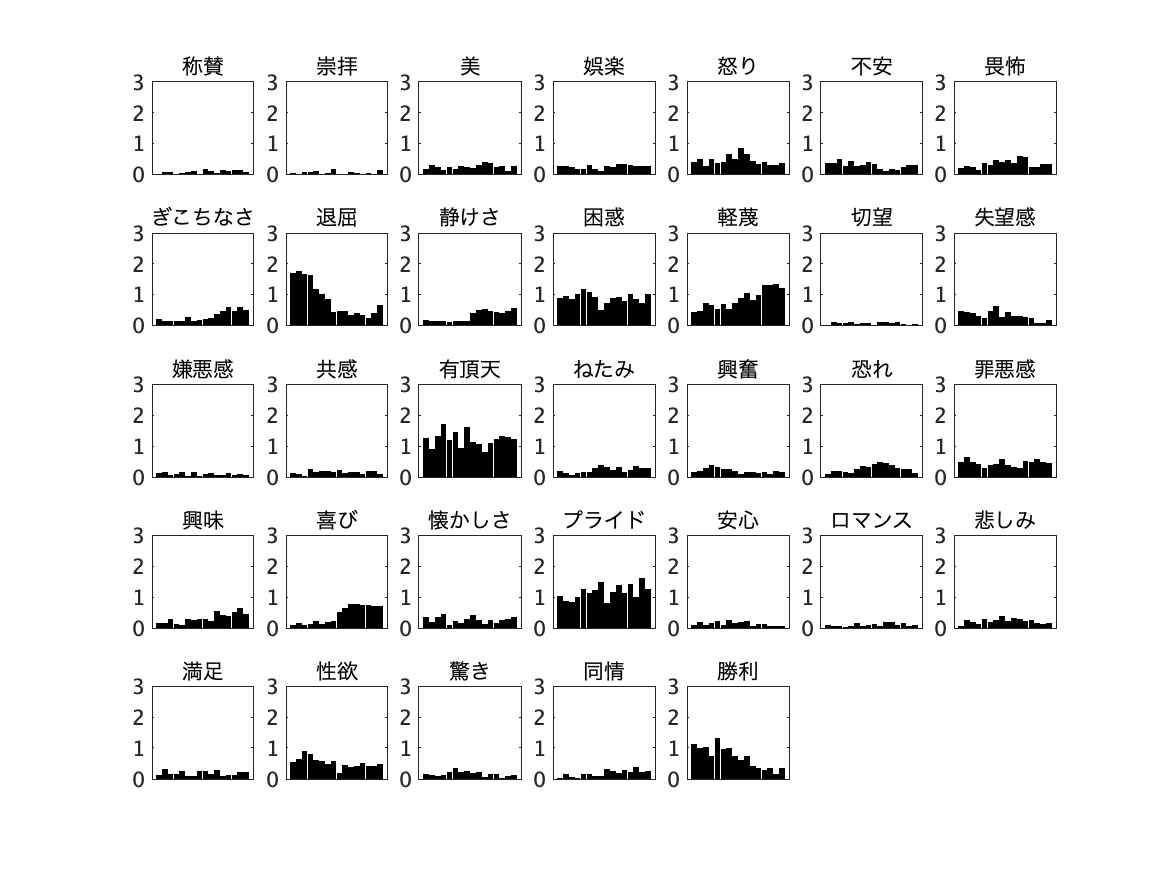

Awkwardness	Boredom	Calmness	Confusion	Contempt	Craving	Disappointment

Admiration	Adoration	Aesthetic	Amusement	Anger	Anxiety	Awe

Disgust	Empathy Entrancement	Envy	Excitement	Fear/Horror	Guilt

Interest	Joy	Nostalgia	Pride	Relief	Romance	Sadness

Satisfaction Sexual desire	Surprise	Sympathy	Triumph

**Externally orientated thinking (EOT) Group**

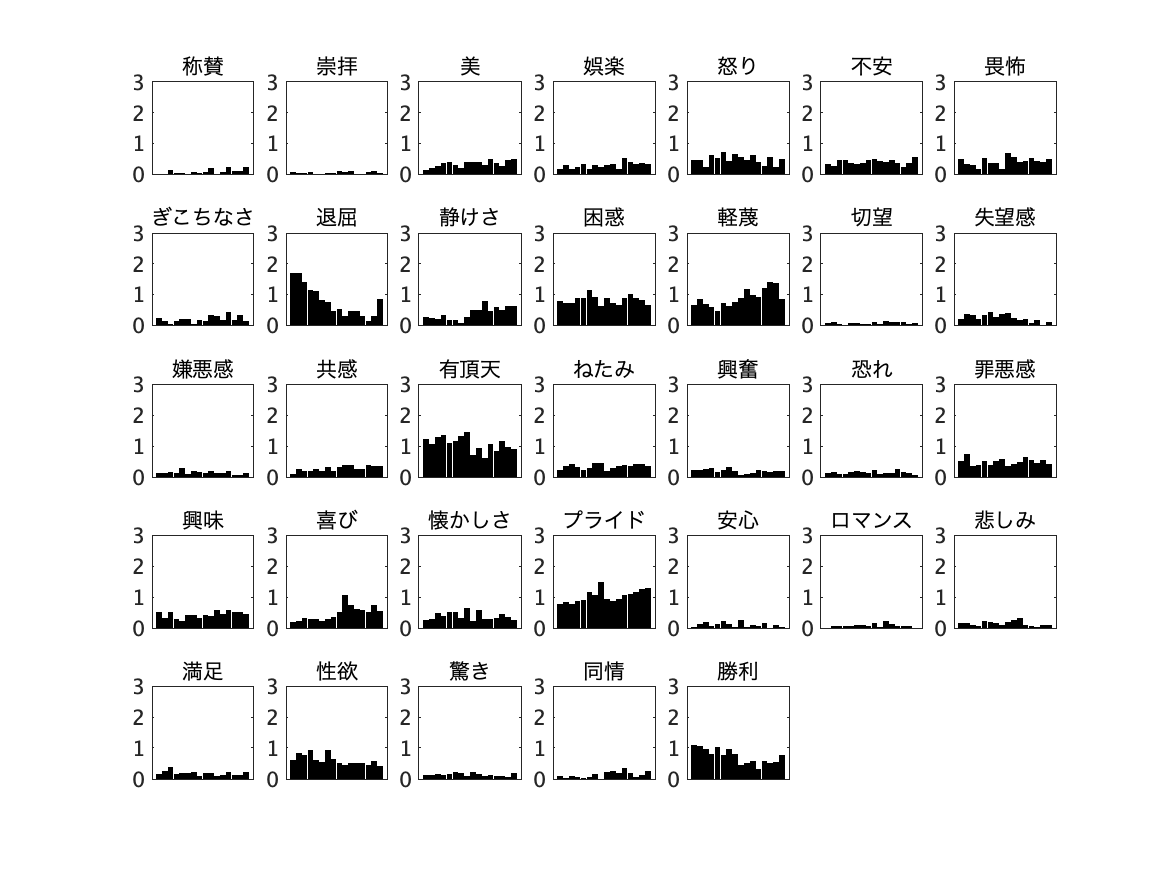

Admiration	Adoration	Aesthetic	Amusement	Anger	Anxiety	Awe

Awkwardness	Boredom	Calmness	Confusion	Contempt	Craving	Disappointment

Disgust	Empathy Entrancement	Envy	Excitement	Fear/Horror	Guilt

Interest	Joy	Nostalgia	Pride	Relief	Romance	Sadness

Satisfaction Sexual desire	Surprise	Sympathy	Triumph

**Non externally orientated thinking (EOT) Group**

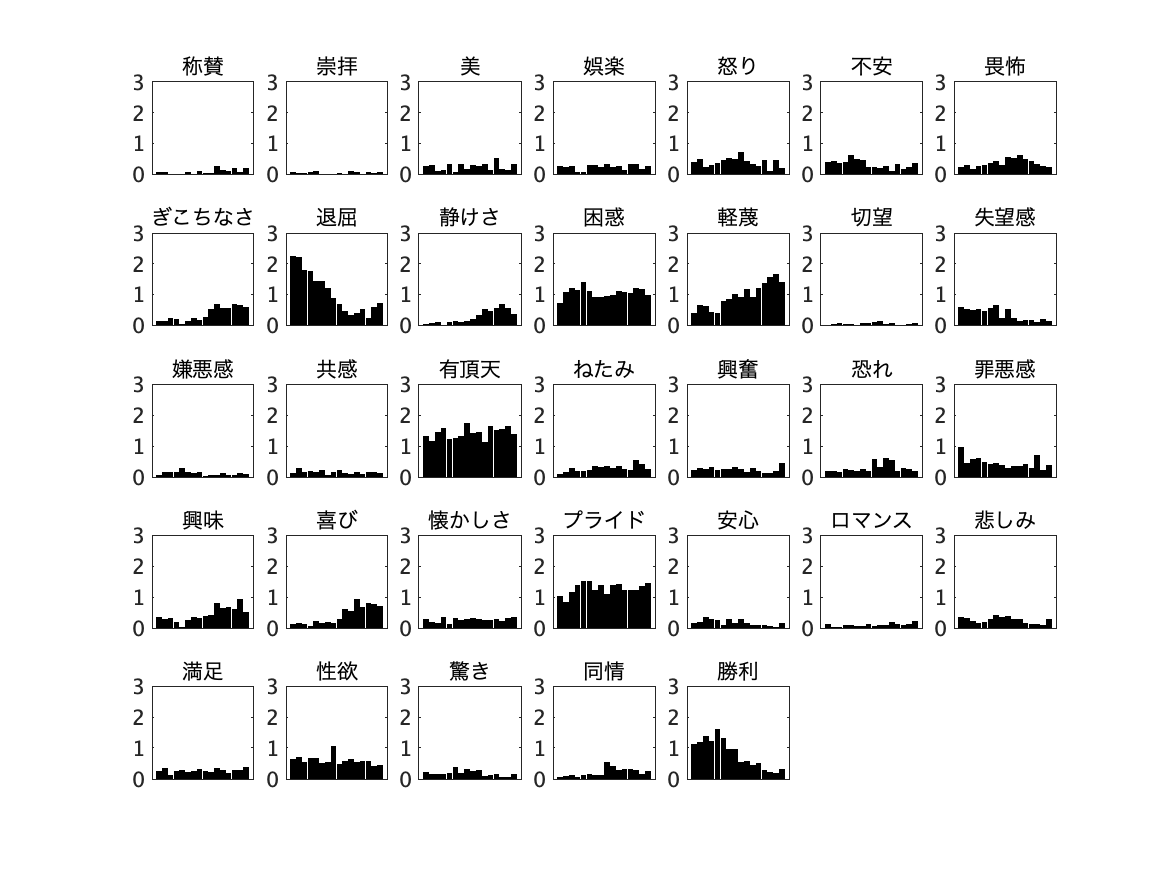

Admiration	Adoration	Aesthetic	Amusement	Anger	Anxiety	Awe

Awkwardness	Boredom	Calmness	Confusion	Contempt	Craving	Disappointment

Satisfaction Sexual desire	Surprise	Sympathy	Triumph

Disgust	Empathy Entrancement	Envy	Excitement	Fear/Horror	Guilt

Interest	Joy	Nostalgia	Pride	Relief	Romance	Sadness

**Figure b. The total number of clicks and the number of different click positions.** The x-axes represent, from left to right, ascending pitches of the 16 notes, while the colours denote Alexithymia (red), depression (blue), and their respective control groups (grey). The total click counts (i.e., the sum of all clicks) and the number of distinct click positions are shown.
